# Notch-signaling is required for mediating between two pattern forming processes during head regeneration in *Hydra* polyps

**DOI:** 10.1101/2024.02.02.578611

**Authors:** Mona Steichele, Lara Sauermann, Qin Pan, Jasmin Moneer, Alexandra de la Porte, Martin Heß, Moritz Mercker, Catharina Strube, Heinrich Flaswinkel, Marcell Jenewein, Angelika Böttger

## Abstract

*Hydra* polyps regenerate lost body parts, including the head. In addition, *Hydra* head tissue has organizer properties thus being able to recruit body column tissue from a host polyp to produce ectopic hydranths after transplantation. These pattern forming processes involve Notch- and Wnt/β-catenin-signaling. *Hydra* head regeneration consists of two parts, hypostome/organizer and tentacle development. Previous work had shown that the Notch inhibitor DAPT blocks hypostome regeneration and organizer formation, but not the appearance of tentacle genes and tentacle tissue. Here we show that the β-catenin inhibitor iCRT14 blocks tentacle regeneration, but not regeneration of hypostome and organizer tissue. Using RT-qPCR gene expression analyzes during head regeneration we found that DAPT inhibits *HyWnt3*- and *HyBMP2/4* expression and expression of transcriptional repressor genes including *CnGsc*, *Sp5* and *HyHes,* while increasing expression of *HyBMP5/8b* and the *c-fos*-related gene *HyKayak.* ICRT14 blocks expression of the tentacle specification factor *HyAlx,* but not expression of *HyWnt3*. Thus, in accordance with regeneration of two head structures we find two signaling and gene expression modules with *HyWnt3* and *HyBMP4* part of a hypostome/organizer module, and *BMP5/8*, *HyAlx* and β-catenin part of a tentacle module. We conclude that Notch functions as an inhibitor of tentacle production to allow regeneration of hypostome/head organizer. Furthermore, with *HyKayak* we present a candidate target gene for HvNotch induced repressor genes. Using siRNA and the Fos/Jun-inhibitor T5224 we show that HyKayak attenuates the expression of *HyWnt3.* Finally, Notch signaling was not required for head regeneration of fresh water polyps of *Craspedacusta*. Polyps of *Craspedacusta* do not have tentacles and thus, after head removal only regenerate a hypostome with a crescent of nematocytes around the mouth opening. This corroborates the idea that Notch-signaling mediates between two pattern forming processes during *Hydra* head regeneration.

## Introduction

The small freshwater polyp *Hydra* is a simple metazoan. It belongs to the pre-bilaterian phylum of cnidaria and consists of a foot, a body column and a head with a hypostome and a ring of tentacles. Asexual reproduction occurs by budding. Sexual reproduction takes place from fertilized eggs when male and female gametes are formed on the *Hydra* body column (reviewed by (Steele, 2012)).

*Hydra* polyps have the capacity for complete regeneration. After being cut into small tissue parts they will regenerate a head and a foot accurately and at the same position as before. This indicates that whole-body pattern information is conserved in the body column during adult life of *Hydra* polyps (reviewed by (Bode, 2003)). Moreover, as observed in 1909 by Ethel Browne, specific *Hydra* tissues, after transplantation into a host polyp, have the capacity to recruit host tissue to form an ectopic head growing out into a whole new hydranth (Browne, 1909; MacWilliams, 1983). These tissues included “peristome at the base of tentacles” and regenerating tips and early buds (according to Ethel Browne). By hypostome-contact grafts it could be shown later, that the tip of the hypostome also produces a signal to induce a second axis growing from the body column of the host by recruiting host tissue. Less “inductive” capacity was found in tissue of the tentacle zone (Broun and Bode, 2002; Mutz, 1930). Embryonic amphibian tissue with such inductive capacity was given the name “organizer” by Hans Spemann and the region where this tissue was taken from was called “center of organization” (Hamburger, 1969; Spemann, 1924). The *Hydra* transplantation phenomena were related to the “organizing” property of transplanted embryonic tissue by (Goetsch, 1926). The “organizer effect” entails a “harmonious interlocking of separate processes which makes up development”, or a side-by-side development of structures independently of each other (Spemann, 1935). In addition to inducing the formation of such structures, the organizer must ensure their patterning (Anderson and Stern, 2016). Formation of new hydranths after transplantation of “organizer” tissue involves the side-by side induction of hypostome tissue and tentacle tissue. Moreover, it includes the establishment of a regularly organized ring of tentacles with the hypostome doming up in the middle. The function of the *Hydra* “center of organization” would then be to pattern hypostome/body column and tentacles and to allow for their harmonious re-formation after head removal.

There is an intriguing similarity in gene expression between the amphibian Spemann organizer and the *Hydra* head organizer (Ding et al., 2017). Spemann-organizers induce a Wnt3-dependent anterior-posterior axis and a BMP-dependent dorsal-ventral axis (Anderson and Stern, 2016). The *Hydra* gene *HyWnt3* is strongly expressed at the hypostome, at the tip of regenerates after head removal, and at the tip of developing buds, all regions that had been indicated to possess inductive capacity in organizer experiments (Broun and Bode, 2002; Browne, 1909; Mutz, 1930)). Additionally, the transcriptional repressor *goosecoid*, is expressed in the dorsal blastopore lip cells of frog embryos and had originally been considered an universal organizer gene (Anderson and Stern, 2016). In the *Hydra* head, *CnGsc*, a *goosecoid-*homolog is prominently (not solely) expressed in head cells between the hypostome and the tentacle zone (Broun et al., 1999), and thus in the organizer tissue as defined by Ethel Browne.

*Hydra* has 11 identified *Wnt* genes, all of which are expressed in the head and/or tentacles. Of those, most are suggested to induce canonical Wnt-signaling through nuclear translocation of β-catenin, while *HyWnt5* and *HyWnt8* have been shown to be associated with non-canonical Wnt-signaling in the planar cell polarity pathway. In addition, most known mammalian BMP-pathway genes have homologs in *Hydra*. These include *Smad*, *HyBMP5/8b* and *HyBMP2/4* (Hobmayer et al., 2001; Lengfeld et al., 2009; Philipp et al., 2009; Reinhardt et al., 2004; Watanabe et al., 2014). Wnt and BMP-pathways have been demonstrated to play a role in *Hydra* regeneration ((Reddy et al., 2020; Reddy et al., 2019) and citations above). After head removal, the expression of *Hy*β*-catenin* and *HyTcf* is upregulated earliest, followed by local activation of *Wnt* genes. Among these, *HyWnt3* and *HyWnt11* appeared within 1.5 h of head removal, followed by *HyWnt1*, *HyWnt9/10c*, *HyWnt16*, and *HyWnt7* (Gufler et al., 2018; Hobmayer et al., 2001; Lengfeld et al., 2009; Philipp et al., 2009; Tursch et al., 2022). Thus *HyWnt3* and *HyWnt9/10* are swiftly induced by injuries. When their activity is sustained organizers can be formed, which induce ectopic heads when the original organizer tissue (the head) is removed (Cazet et al., 2021; Tursch et al., 2022). Recently, a Wnt3/β-catenin/Sp5 feedback loop was suggested to be involved in *Hydra* head patterning (Moneer et al., 2021; Münder et al., 2013; Vogg et al., 2019).

The expression patterns of *Wnt*- and *BMP*-genes can be interpreted as an indication of tentacles, buds and the main body axis of the polyps being repetitive structures expressing *Wnt*-genes at the apical end and *BMP5/8b* at the basal end (Meinhardt, 2012; Pan et al., 2024). These could set up opposing signaling gradients to pattern the *Hydra* body axis and possibly also the bud and tentacle axes. The bud expresses *HyWnt2* and later *HyWnt3* at the tip and *BMP5/8b* at the base. The tentacles also express *HyBMP5/8b* at the base and *HyWnt5* at the tip. As Hans Meinhard pointed out, in evolutionary terms the tentacles may therefore be considered as colonialized buds (Meinhardt, 2012). In any case, tentacles and hypostome can be interpreted as independent structures.

In addition, our previous investigations had revealed that the Notch-pathway was instrumental for head regeneration and organizer formation by supporting the expression of a strong *HyWnt3*-signal in regenerating head tissue. Notch inhibition with the presenilin inhibitor DAPT or the NICD-inhibitor SAHM-1 prevented head regeneration and blocked *HyWnt3* expression in regenerates, while not preventing expression of the tentacle boundary gene *HyAlx* and the tentacle metalloprotease gene *HMMP*. However, the latter did not obtain their correct expression patterns and thus proper tentacles were not formed. Similar experiments using a transgenic *Hydra* strain expressing a HvNotch-hairpin RNA confirmed the regeneration phenotypes seen with pharmacological inhibitors (Pan et al., 2024). Strikingly, transplantation experiments had revealed that DAPT-treated regenerating head tissue had lost the capacity to form an organizer (Münder et al., 2010; Münder et al., 2013).

Here we have further investigated the role of Notch-signaling during apical head regeneration. We compared the effect of the Notch-inhibitor DAPT with the effect of the β-catenin inhibitor iCRT14 (Gonsalves et al., 2011; Gufler et al., 2018). While, similar to DAPT, iCRT14-treated animals did not regenerate complete heads, the expression of *HyWnt3* at the regenerated hypostome was not blocked. Accordingly, iCRT14-treated - in contrast to DAPT-treated - regenerating tips retained the ability to form a second axis when transplanted into the body column of an untreated host animal. We also investigated the effect of these inhibitors on the gene expression dynamics of *HyWnt-* and *HyBMP*-genes and transcriptional regulators *Hydra Sp5*, *HyAlx, HyHes* and *CnGsc* during *Hydra* head regeneration by RT-qPCR. Our results clearly reveal that sustained expression of *HyWnt3* and hypostome/organizer formation after head removal are controlled by Notch-signaling, and not by β-catenin activity. In contrast, expression of the tentacle specification gene *HyAlx* and formation of tentacles are dependent on β-catenin activity. Additionally, we noted that Notch-inhibition increases the expression of *HyBMP5/8b*, a gene primarily expressed at tentacle boundaries, while blocking the expression of *HyBMP2/4*, a gene expressed in the head and body column. Moreover, Notch was required for inhibition of the *c-fos* homolog-*HyKayak,* which we suggest to be a negative regulator of *HyWnt3* and a likely candidate for a target of Notch regulated transcriptional repressors.

We conclude that Notch activity functions in head regeneration to mediate between two independent patterning systems comprising hypostome- and tentacle regeneration. In apical regenerates this probably works through inhibition of the tentacle system in a spatially and temporarily regulated manner. It involves Notch-mediated inhibition of *HyBMP5/8b* and direct or indirect activation of *HyWnt3* and *HyBMP2/4* expression.

## Methods

### Animal treatment

*Hydra* polyps were cultured in *Hydra* Medium (HM) (0.29mM CaCl_2_, 0.59 mM MgSO_4_, 0.5 mM NaHCO_3_, 0.08 mM K_2_CO_3_ dissolved in MiliQ water) at a constant temperature of 18°C. They were fed with freshly hatched *Artemia nauplii* 2-3 times per week, with the exception of two days prior to conducting the experiments. For regeneration experiments, all animals were decapitated at 80% of their body length and left to regenerate for 2 days in HM containing the respective inhibitors dissolved in 1% DMSO. Control animals were left to regenerate in HM with 1% DMSO. Treatments included 35 µM DAPT/1% DMSO, 5 µM iCRT14/1% DMSO or 7.5 µM T5524 for 8 hrs, 24 hrs, 36 hrs, 48 hrs after head removal. Time point 0 refers to animals immediately after the head was cut off. The inhibitor/DMSO containing medium was renewed every 12-14 hrs.

*Craspedacusta sowerbii* polyps were grown in modified HM (0.29mM CaCl_2_, 0.59 mM MgSO_4_, 0.5 mM NaHCO_3_, 0.08 mM K_2_CO_3_ dissolved in MiliQ water) at 19°C. They were fed with *Brachionus calyciflorus* twice a week. For regeneration experiments, all animals were decapitated at 80% of their body length and left to regenerate for 3-4 days in HM containing the respective inhibitors dissolved in 1% DMSO. Control animals were left to regenerate in HM with 1% DMSO. Treatments included 35 µM DAPT/1% DMSO or 5 µM iCRT14/1% DMSO for 8 hrs, 24 hrs, 36 hrs, 48 hrs, 72 hrs or 96 hrs after head removal. Time point 0 refers to animals immediately after the head was cut off. The inhibitor/DMSO containing medium was renewed every 12-14 hrs.

### Standardising conditions for RTqPCR

For quantitative estimates of gene expression dynamics during *Hydra* head regeneration over time we performed real-time quantitative reverse transcription (RT)/PCR (qPCR) experiments. We used a fluorescence-based qPCR-method and adhered to the quality standards of the MIQE guidelines (Bustin et al., 2009). After *in silico* primer design, each primer pair was empirically validated for (1) specificity defined by a single melt peak corresponding to a unique band of expected size, (2) efficiency defined by doubling of the signal in every cycle and (3) sensitivity defined by a broad linear range, and reproducibility. Primers and gene accession numbers are listed in table S1. Total RNA was isolated from *Hydra* polyps and RNA quality was tested with the Agilent bioanalyzer. Only RNA with an integrity number higher than 8 was used for cDNA synthesis. During head regeneration, mRNA for RT-qPCR was isolated from whole regenerates collected after 8, 24, 36 and 48 hrs (t= 8, 24, 36, 48). Immediately after head removal the sample for t=0 was obtained. All experiments included three biological replicates with three technical replicates each. Quantitative gene expression for each gene was calculated as the ratio of target gene expression to housekeeping gene average (relative normalized gene expression). We plotted the relative normalized gene expression of analyzed genes against the regeneration time points. Housekeeping genes included *GAPDH, PPIB, RPL13*.

### Regression analysis of comparative expression levels

To visualise temporal changes in expression levels of different genes, we used appropriate regression methods. In particular, we used generalised additive models (GAMs) (Hastie & Tibshirani, 1990; Wood, 2017) enabling the visualisation of nonlinear dependencies on the time-dependent variables based on appropriate regression splines (Wood, 2017). Here, we used the Tweedie probability distribution (Kokonendji, Demetrio, et al., 2004; Kokonendji, Dossou-Gbete, et al., 2004) which is known to describe non-negative (possibly over dispersed) data well – in particular if mean values are close to zero. Temporal autocorrelation of model residuals has been investigated based on pacf-plots (Field et al., 2012; Wood, 2017) and was not apparent. The optimal amount of smoothness of regression splines has been estimated separately for each temporal expression pattern based on generalized cross-validation methods (Wood, 2017). For the analysis of expression patterns relative to the control (DMSO) type, the response variable in regression analysis has been defined by dividing separately for each experiment/time point the mean value of the repeated measurements of the treatment of interest (DAPT respectively iCRT) by the mean value of the repeated measurements of the corresponding DMSO treatment from the same experiment/time point.

### Semithin sections with Richardson staining

Animals were fixed with 4% PFA and prepared for semithin sectioning by re-fixation in 1% osmium tetroxide solution for 2 hours. Samples were washed with water and dehydrated 4 times with serial acetone dilutions (30%, 50%, 70%, 90%, four times 100%). Finally, they were embedded in Spurr low viscosity embedding medium standard mix, which was exchanged 4 times, and dried after each exchange for 24 hours at 60°C in a cuboid shape. The resin embedded probes were sectioned with a semi-diamond and stained after Richardson on a microscope slide. One drop of colour solution (1% Azur in H_2_O and 1% methylene blue in 1% Na_2_B_4_O_2_ in H_2_O mixed 1:1) covering the semithin sections was heated to 80°C for 30 seconds and cleansed with water. After drying, the slides were analysed with a brightfield microscope.

### Histochemistry of polyps

Polyps were relaxed in 2% urethane and fixed with 4% PFA in HM for 1 hr. They were permeabilized with ice-cold 100% ethanol and blocked in 0.1% Triton/1% BSA in PBS. For phalloidin staining they were incubated with Phalloidin-iFluor 488 (abcam, ab176753) (1:500) for 1 hr, followed by DAPI (1:1000) staining before mounting on slides with Vectashield. Slides were analysed with a Leica SP5 point scanning laser confocal microscope equipped with oil immersion HCX PL APO Lambda Blue 20x 0.7 and 63x 1.4 objective lenses. Alexa-488 fluorochromes were visualized with an argon laser at an excitation wavelength of 488 nm and emission filters of 520-540 nm and a diode laser at an excitation wavelength of 405 nm and emission filter at 450-470 nm was used for DAPI. The produced light optical serial sections were stacked with the ImageJ plugin StackGroom to produce 3D images of the treated polyps. DAPI staining of nematocyte capsules was done according to (Szczepanek et al., 2002).

### Fluorescent in situ hybridization

This experiment was carried out as previously described (Siebert et al., 2019).

### Transplantation experiments

Non-budding *Hydra* polyps were pre-treated with 5 µM iCRT14/1% DMSO in HM for 24 hrs. After that they were bisected at 80% of the body column underneath the head and left to regenerate in iCRT14 treated HM for another 24 hrs. The newly regenerated head region (top20%) was grafted onto a blue host animal (treated with Ewan’s Blue for two weeks) at about 50% of the body column. After 3 hrs the rod was removed and the animals left in HM for another 48 hrs. Finally, the animals were classified for the presence of newly formed secondary axes displaying a clear hypostome and tentacles. Tissue recruitment was recognized by the blue/white colour distribution within the new axes.

### ShRNA knockdown

shRNA design and production were done according to Karabulut’s protocol (Karabulut et al., 2019). For electroporation, 30 budless Hydra polyps were washed 5 times with MiliQ water and incubated for 45 min in MiliQ water. Then, excess water was removed and replaced with 200 μl of a 10 mM HEPES solution at pH 7.0. The suspended animals were then transferred into a 4mM gap electroporation cuvette, and 4 μM of purified shRNA or scramble shRNA was added to the cuvette. The mixture was mixed by gently tapping the cuvette for 5 times and incubated for 5 min to let animals relax before electroporation. The electroporation was carried out using the BTX Electro Cell Manipulator 600 by setting up the condition to 250 V, 25 ms, 1 pluse, 125 μF capacitance. 500 μl of restoration medium (80 % HM and 20% dissociation medium (3.6 mM KCl; 6 mM CaCl_2_; 1.2 mM MgSO_4_; 6 mM sodium citrate; 6 mM sodium pyruvate; 6 mM glucose; 12.5 mM TES; and 50 mg/ml rifampicin, pH6.9)) was added into the cuvette immediately after electroporation. The entire volume of eletroporated animals were then transferred into a petri dish. In our experiment, three times of electroporation were done every two days to achieve a significant knockdown of HyKayak. And two hairpins of Kayak were used for electroporation at 1:1.

### Monoclonal anti-HyKayak-antibody

Mice were immunized with fusion protein Hydra_KAYAK-HIS (amino acid of HyKAYAK: 1-111) using a mixture of 50µg protein, 12 µl Oligo 1668 (500 pmol/µl) and 150 ul IFA in a total volume of 400 µl. After > six weeks a single boost was given with the same mixture except for the IFA, which was omitted. Fusion with Ag8 myeloma cells was performed using standard procedures. Candidate selection was based on positive selection using KAYAK-HIS and negative selection using *Hydra*_HES-HIS. Hybridoma kayak 3C10-1-1 and 13A4-1-1 were cloned using standard procedures and subsequently grown for antibody production.

### Multiple sequence alignment and phylogenetic analysis

The multiple sequence alignment was done using Clustal Omega. The conserved domains were identified by PROSITE. The phylogenetic trees were produced by MEGA. The protein sequences for comparison were retrieved from Uniprot and NCBI.

### Subcellular fractionation and Western blot

500 Hydras were dissociated into single cells with 10 mL dissociation medium by pipetting. After centrifuge at 2000 g for 10 min, the cellular pellet was resuspended in 500 μl RIPA buffer (25 mM Tris HCl, pH 7.5, 150 mM NaCl, 1 % NP40, 1 % sodium deoxycholate, 0.1 % SDS, 10 ng/ml Pepstatin A, 10 ng/ml Aprotinin, 10 ng/ml Leupeptin, 0,5 mg/ml Pefablock) and incubated for 20 min on ice. Subsequently, the mixture was homogenized with a Dounce homogenizer for 30 times and then centrifuged for 10 min at 1000 g. The resulting supernatant including cytoplasmic proteins was collected and labelled as CP. The pellet was treated with 500 μl RIPA buffer, and then sonicated at 180 watts for 3 min (in rounds of 10 sec sonication and 50 sec rest for each cycle). After centrifuging at 14000 g for 30 min, the supernatant was collected and labelled as nuclear proteins (NP), the pellet was resuspended with the same volume of RIPA buffer and kept for SDS-PAGE gel analysis.

For DNase treatment, the pellet from the second centrifuge was resuspended with 500 μl RIPA buffer supplemented with 200 units/mL DNase, 10 mM CaCl_2_ and 10mM MgCl_2_, and incubated at room temperature for 15 min. After centrifuging at 1000 g for 10 min, the supernatant was collected, while the pellet was resuspended in 500 μl RIPA buffer with 2 M NaCl and incubated on ice for 10 min. Then the same centrifuge was done and the supernatant and pellet was collected for gel analysis.

Western blots were stained with the inhouse mouse anti-Kayak monoclonal antibody.

### Co-immunoprecipitation

HEK293T cells were transferred with C-terminal HA-tagged Kayak and N-terminal GFP-tagged Jun-epi or Kayak using Lipofectamine 2000 (11668030, Thermo Fisher). The GFP-trap agarose beads (ABIN509397, ChromoTek) were used for immunoprecipitation as described previously (Heim et al., 2014; Webby et al., 2009). Western blot was stained with the following primary antibodies: mouse anti-GFP antibody (11814460001, Roche) and rabbit anti-HA antibody (H6908, Sigma Aldrich).

### Identification of *Craspedacusta* genes

*Craspedacusta* totalRNA was extracted from 120 polyps using the Qiagen RNeasy Mini Kit. RNA-quality was verified with the Agilent bioanalyzer, the RNA was then transcribed into cDNA and cDNA was sequenced with Illumina. The resulting gene sequences were aligned and by comparison with sequences for *HyWnt3*, *NOWA*, *HyAlx* and *Sp5* the corresponding *Craspedacusta* cDNA-sequences could be identified (*CsWnt3*, *CsNOWA*, *CsAlx* and *CsSp5*) and confirmed by sequencing of cDNA-clones obtained after RT-qPCR from *Craspedacusta* total RNA.

## Results

## 1. Hypostome formation in iCRT-treated, but not in DAPT-treated regenerates

*Hydra* polyps treated either with iCRT14 as described by Cazet 2021 and Gufler 2018 (Cazet et al., 2021; Gufler et al., 2018), or with the Notch-inhibitor DAPT, as described by Münder 2013 (Münder et al., 2013), fail to regenerate a complete head after decapitation. DAPT blocks Notch intramembrane proteolysis regulated by presenilin and prevents NICD-translocation to the nucleus, thus phenocopying loss of Notch function in several organisms including *Hydra* (Dovey et al., 2001; Geling et al., 2002; Käsbauer et al., 2007; Micchelli et al., 2003; Pan et al., 2024). iCRT14 inhibits the interaction of nuclear β-catenin with TCF in mammalian cell lines and in *Hydra* (Gonsalves et al., 2011; Gufler et al., 2018).

First, we treated *Hydra* polyps with 5 µM iCRT14 for 48 hrs after head removal, and observed that they did not regenerate their heads during the time of treatment, while control animals, treated with 1% DMSO (the solvent for iCRT14 and DAPT), clearly showed regularly spaced tentacle buds at this time point (Fig. 1A). iCRT14 and DMSO were then replaced with normal *Hydra* medium. Control animals regenerated heads with long tentacles 24 hrs later (72hrs), however iCRT14-treated animals did still not show tentacle buds up to 48 hrs after iCRT14 removal (96 hrs). For comparison, treatment of head regenerates with DAPT had revealed in our previous study that proper heads could also not be regenerated during the time of treatment. When DAPT was then removed from the medium, irregular heads, dominated by tentacle tissue, developed in 20 % of regenerates (Münder et al., 2013).

**Figure 1:**
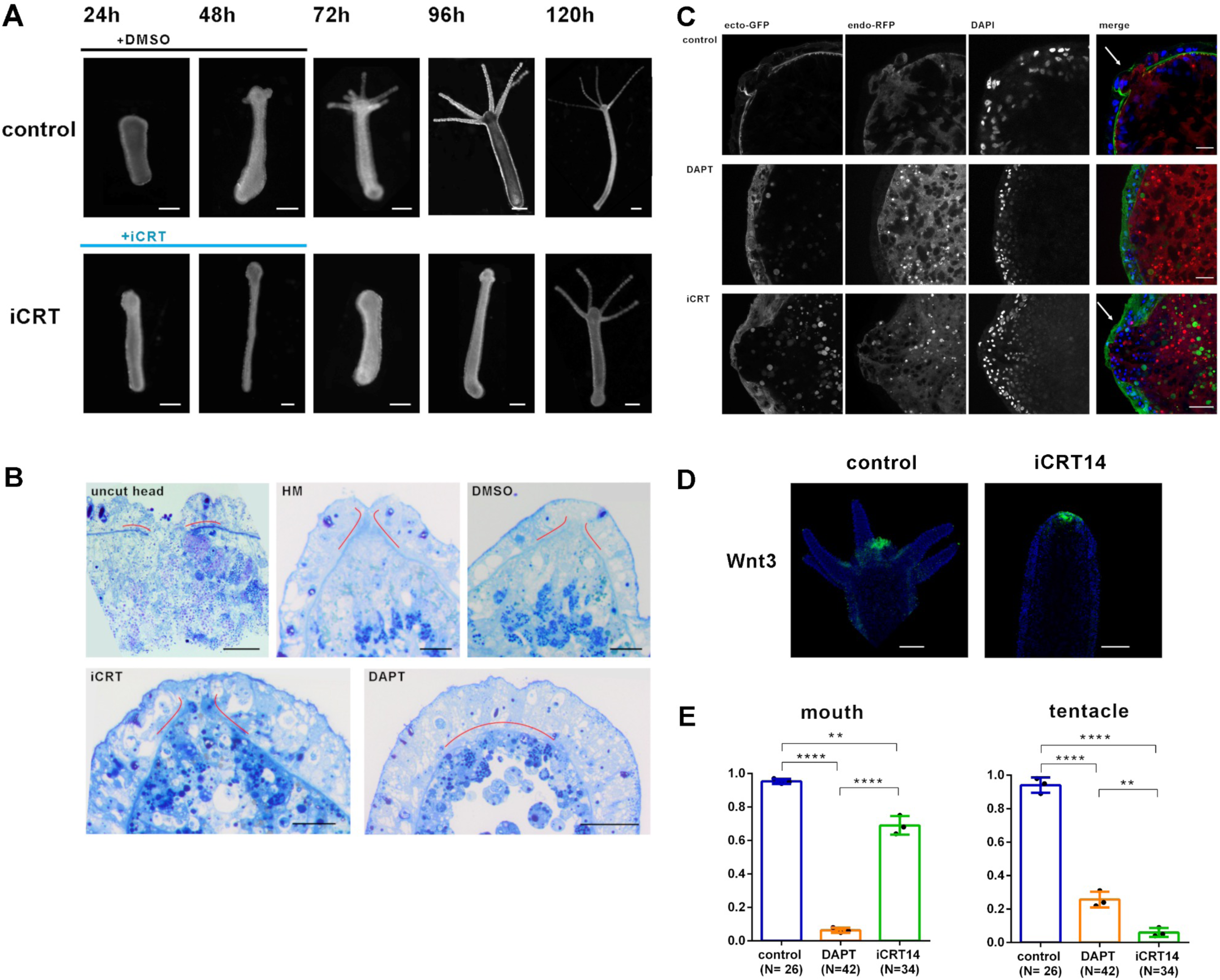
Regeneration of *Hydra* head structures. **(A)** Head regeneration progress of *Hydra* polyps at indicated time points after head removal in control - or iCRT14-medium. **(B)** Semi-thin sections covering the hypostome region of *Hydra* polyps after Richardson-staining: untreated polyp (uncut head); polyps 48 hrs after head removal and regeneration in *Hydra* medium (HM) and HM with DMSO (control), DAPT or iCRT14. Mesoglea appears as dark blue line, red lines are added to highlight the mesoglea at the hypostome region. Scale bar: 20 µm. **(C)** Confocal stack images covering the regenerated hypostome region of *Hydra* polyps of strain AEP ‘Water melon’ 48 hrs after head removal and regeneration in HM with DMSO (control), DAPT or iCRT14; GFP present in ectodermal cells, dsRed present in endodermal cells and DAPI-DNA-stain are imaged as indicated. Right hand panels show merged images. White arrows indicate hypostomal opening in iCRT-treated and control animals, but not in DAPT-treated polyps. Scale bar: 10 µm. **(D)** Fluorescent In-situ hybridization for *HyWnt3* (green) expression in polyps 48 hrs after head removal and regeneration in iCRT14 or in DMSO control, as indicated. Scale bar: 100 µm, DAPI in blue. **(E)** Quantification of regeneration of *Hydra* head structures: mouth and tentacles, 48 hrs after head removal and regeneration in DMSO control, DAPT and iCRT14. Data shown as mean ± SEM, *: p=0.05, **: p=0.01, ***: p=0.001, ****: p=0.0001.

To further inspect the morphology of head regenerates treated with DAPT or iCRT14, semithin sections were prepared 48 hrs after head removal and histologically stained with Richardson tissue stain. Amongst other structures, this dye stained the mesoglea dark blue. Fig. 1B shows middle sections of polyps. The mesoglea is emphasized by red lines. The hypostome of the polyp can be recognized by a ‘gap’ in the mesoglea. After head removal, the hypostome is regenerated in polyps treated with DMSO and iCRT14, but not with DAPT. Head regeneration of ‘water melon’ AEP-vulgaris *hydra* polyps showed a similar result (Fig. 1C). These polyps express green fluorescent protein (GFP) in the whole of the ectoderm and red fluorescent protein (RFP, dsRed) in the whole of the endoderm (polyps were a kind gift from Rob Steele, UC Irvine). Fig. 1C shows optical middle sections obtained by laser scanning microscopy clearly representing a mouth opening. Again, hypostome morphology is recovered in animals after regeneration in DMSO and iCRT14, but not in DAPT. Quantification of regenerated hypostomes and tentacles in DAPT- and iCRT14-treated regenerates in comparison with control animals revealed that 70 % of iCRT14-treated animals regenerated an intact hypostome with a detectable mouth opening, whereas tentacles were not formed (Fig. 1E). In contrast, DAPT-treated animals did not regenerate a mouth opening, and in 25% of regenerates aberrant tentacles were observed at the tips of regenerates, as previously described (Münder et al., 2013). The apparent regeneration of a hypostomal mouth opening in iCRT14-treated polyps prompted us to perform fluorescent in situ hybridisation for *HyWnt3* in such regenerates. As shown in Fig. 1D, hypostomal *HyWnt3* expression was evident in control regenerates and showed a very similar pattern in regenerates treated with iCRT14. This was different from DAPT-treated regenerates, which do not express *HyWnt3* (Münder et al., 2013).

## 2. Organizer formation observed in iCRT14-treated regenerates

Previously, we had shown that DAPT-treated regenerating *Hydra* heads lacked organizer activity, as they did not induce the formation of ectopic hydranths when transplanted into the body column of a host animal (Münder et al., 2013). This was in accordance with the loss of *HyWnt3* expression in Notch-inhibited regenerates. We now asked the question whether iCRT14-treated head regenerates would retain organizer properties, as they do express *HyWnt3*. We therefore transplanted regenerating *Hydra* heads (upper 20% of polyps) 24 hrs after head removal and treatment with iCRT14 or DMSO (for control) into the body column of Ewan’s blue stained host animals. Fig. 2B shows that 80 % of control regenerates formed ectopic hydranths after transplantation into the body column of the host. However, 80 % of iCRT14-treated regenerates were also able to form ectopic hydranths and most of them recruited host tissue, indicating organizer activity. This is in accordance with their expression of *HyWnt3*. From these and previous data we conclude (1) organizer activity correlates with the presence of *HyWnt3* expression; (2) activation of *HyWnt3* during the regeneration process is not dependent on β-catenin transcriptional activity and (3) *HyWnt3* must signal via a non-canonical Wnt-signaling pathway in iCRT14 treated regenerates.

**Figure 2:**
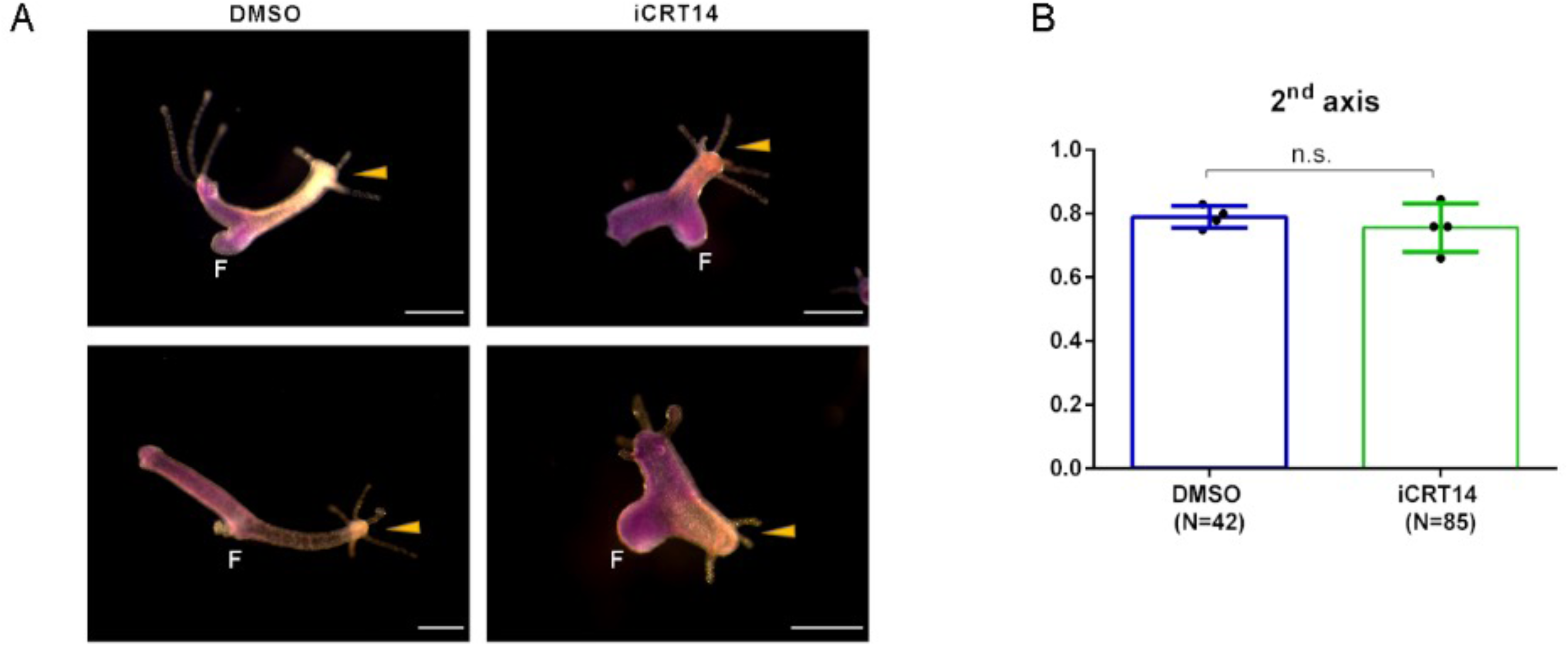
Organizer activity of regenerating *Hydra* head tissue. Tissue from regenerating tip of polyps 24 hrs after head removal in HM with DMSO (control), or iCRT14 was transplanted into the middle of blue host polyps stained with Ewan’s blue. **(A)** Microscopical images were taken 48 hrs after transplantation. Newly formed second axes are indicated by yellow arrows. The transplanted tissue appears orange, the host tissue appears blue, feet are indicated by F. **(B)** Percentage of transplants forming new axes in HM with DMSO (control), or iCRT14; differences are not significant (n.s.)

## 3. Comparison of gene expression dynamics during *Hydra* head regeneration in DAPT-treated and iCRT14-treated animals

In order to follow the recovery of head specific gene expression after head removal we conducted RT-qPCR analyzes from tissue that was left to regenerate. We compared gene expression in regenerates treated with DAPT or with iCRT14, both compounds were administered with 1% DMSO in HM. For control, the polyps were treated with 1% DMSO in HM without additional compounds.

### 3.1. Effect of Notch inhibition on gene expression dynamics during head regeneration in Hydra

In a previous transcriptome analysis of DAPT-treated *Hydra* polyps, beside *HyHes*, the tentacle boundary gene *HyAlx,* the “organizer” gene *CnGsc* and the *Hydra Sp5* gene had been suggested to be potential direct Notch-target genes (Moneer et al., 2021). The same analysis had revealed that the fos-related transcription factor *HyKayak* was up-regulated when Notch-signaling was blocked.

We now performed RT-qPCR analysis to compare gene expression dynamics of these genes during head regeneration 0, 8, 24, 36 and 48 hrs after head removal. Animals were either treated with 30 µM DAPT in 1% DMSO, or 1% DMSO as control. Timepoint 0 was measured immediately after head removal. The results of these analyses revealed that *HyHes* expression was clearly inhibited by DAPT during the first 36 hrs after head removal (Fig. 3A), confirming previously published data which had indicated HyHes as a direct target for NICD (Münder et al., 2010). *HyAlx* expression levels were slightly up-regulated after 24 hrs, but later partially inhibited by DAPT (Fig. 3B). *CnGsc* expression under DAPT treatment initially (8hrs) was comparable to control levels, but then it was strongly inhibited (Fig. 3C). This corresponds with the observed absence of organizer activity in regenerating *Hydra* tips (Münder et al., 2013). Interestingly, a similar result was seen for *HySp5* expression, which was also normal at 8 hrs but was then inhibited by DAPT at later time points (Fig. 3D). *HyKayak*, while not affected after 8 hrs, was strongly overexpressed between 24 and 36 hrs of regeneration in DAPT-treated polyps in comparison to control regenerates (Fig. 3E). However, at the 48 hr time point expression appeared normal.

**Figure 3:**
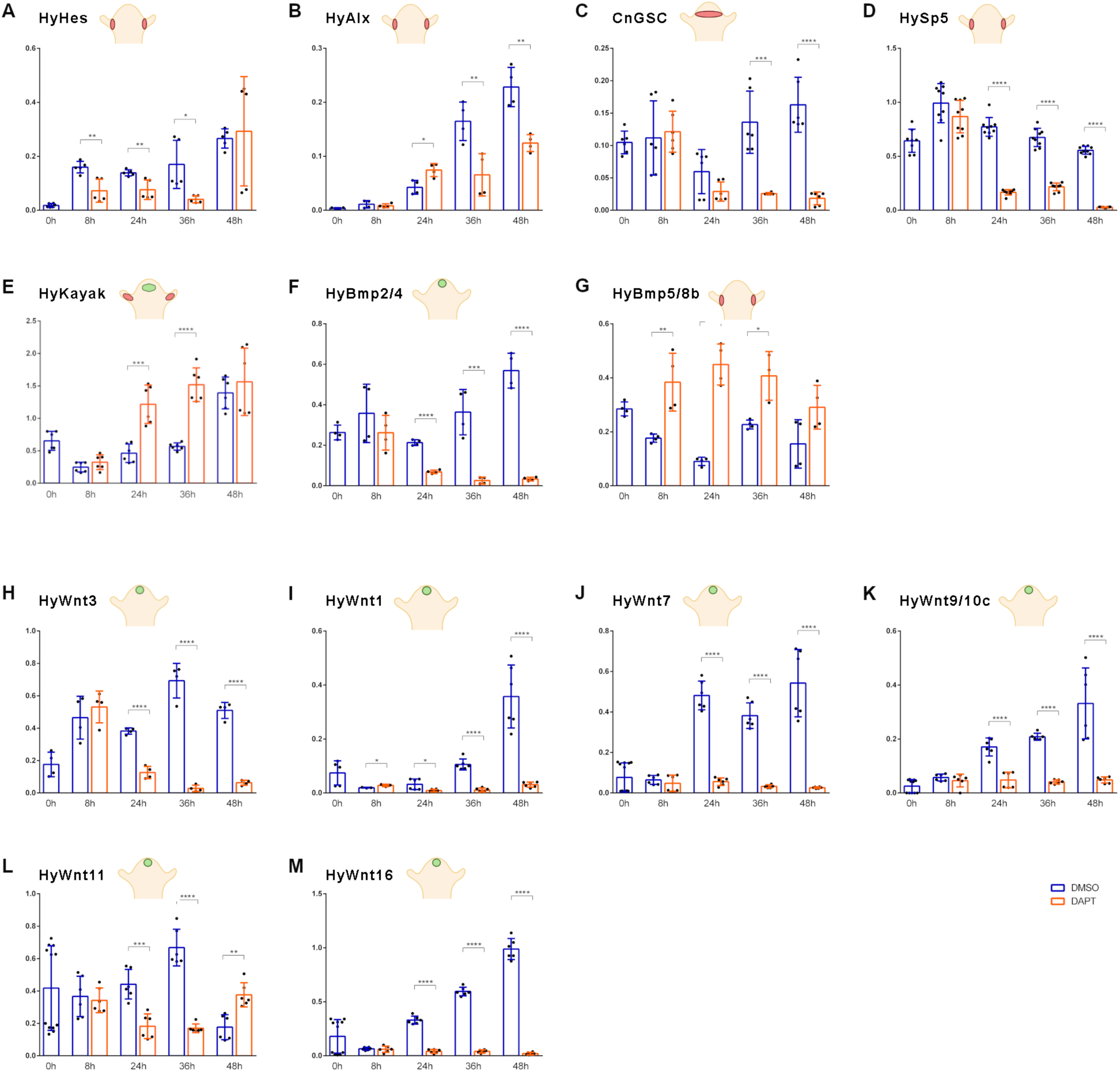
Gene expression dynamics of selected genes in DAPT inhibited regenerates. RT-qPCR measurements quantifying gene expression of **(A)** *HyHes* **(B)** *HyAlx* **(C)** *Goosecoid* **(D)** *Sp5* **(E)** *HyKayak* **(F)** *BMP2/4* **(G)** *BMP5/8b.* **(H-M)** *Wnt3, Wnt1, 7, 9/10c, 11 and 16* during 48 hrs of *Hydra* head regeneration in HM with DAPT (orange) or DMSO (control), (blue). *Hydra* cartoons indicate gene expression patterns according to published in situ hybridisation data and single-cell sequencing atlas (Broun, Sokol et al. 1999, Hobmayer, Rentzsch et al. 2000, Smith, Gee et al. 2000, Reinhardt, Broun et al. 2004, Lengfeld, Watanabe et al. 2009, Münder, Käsbauer et al. 2010, Watanabe, Schmidt et al. 2014, Siebert, Farrell et al. 2019, Vogg, Beccari et al. 2019); Relative normalized expression was calculated against the housekeeping genes *GAPDH*, *RPL13*, *EF1alpha* and *PPIB*. Regeneration time is shown on x-axes; t=0 refers to animals immediately after the head was removed. Data is shown as mean ± SEM, *: p=0.05, **: p=0.01, ***: p=0.001, ****: p=0.0001.

In addition, we tested the expression dynamics of the two *BMP*-homologs described in *Hydra*, *HyBMP5/8b* and *HyBMP2/4.* They have exclusive expression patterns in the head. *BMP2/4* is expressed in endodermal and ectodermal epithelia cells of the head, while *BMP5/8b* expression is restricted to the base of tentacles and is not found in apical head cells (Reinhardt et al., 2004; Siebert et al., 2019; Watanabe et al., 2014). Interestingly, the two *BMP*-genes were conversely affected by Notch inhibition. *HyBMP2/4* expression was blocked with DAPT beginning at 24 hrs of regeneration (Fig. 3F). In contrast, *HyBMP5/8b* expression was drastically increased (Fig. 3G).

We had previously shown by in-situ hybridization that *HyWnt3* is not expressed in DAPT-treated head regenerates (Münder et al., 2013). This was confirmed now by RT-qPCR measurements, which revealed that *HyWnt3* expression was comparable to the control group 8 hrs after head removal. However, after this time point, its expression was strongly inhibited by DAPT and almost completely lost after 36 and 48 hrs (Fig. 3H). Eventually we analyzed most of the *Wnt*-genes suggested to engage in canonical Wnt-signaling, including *HyWnt 1*, 7, 9/10c, 11 and 16 (Lengfeld et al., 2009). In the presence of DAPT, these genes all exhibited similar expression levels to the control group 8 hrs after head removal, but between 24 and 48 hrs the expression of *HyWnt 1*, *7*, *9/10* and *16* declined to almost zero (Fig. 3I-M). As an exception, *HyWnt11* was only partially inhibited and even appeared up-regulated after 48 hrs (Fig. 3L).

In summary, RT-qPCR analyzes showed that Notch-signaling during *Hydra* head regeneration is necessary for activating all *HyWnt*-genes, which are expressed in the *Hydra* head region and implicated in canonical Wnt-signaling. Notch is also necessary for activation of *BMP2/4* expression, a gene expressed in the *Hydra* head and body column. Moreover, Notch is contributing to the expression of transcriptional repressor genes, *HyHes* and *CnGsc*. In contrast, *HyBMP5/8b* and *HyKayak* seem to be subject to inhibition by Notch-signaling. *HyAlx*, although previously identified as a Notch-target gene, is only partially inhibited by DAPT during head regeneration.

### 3.2. Effect of β-catenin inhibition on gene expression dynamics during head regeneration in Hydra

Next, following the same procedure as described for DAPT, we compared the gene expression dynamics of iCRT14-treated regenerates with control regenerates. We found that the expression the Notch-target gene *HyHes* remained similar to control regenerates up to 24 hrs, but then was attenuated (Fig. 4A), possibly due to failure of tentacle boundary formation, the tissue where *HyHes* is strongly expressed. *HyAlx* expression was completely abolished by iCRT14 consistent with the observation that iCRT14-treated head regenerates did not regenerate any tentacles (Fig. 4B). Furthermore, we found that *CnGsc* levels in iCRT14 treated regenerates remained similar to control regenerates up to 24 hrs, but reached only half of the control levels after 48 hrs, similar to *HyHes* (Fig. 4C). Sp5 did not significantly respond to iCRT14 treatment (Fig. 4D). The expression of *HyKayak* was decreased at 8 hrs after head removal in the presence of iCRT14, came back to normal after 36 hrs and was suddenly increased after 48 hrs (Fig. 4E), correlating with inhibition of the *HyHes* repressor. There were no significant changes in the expression dynamics of *HyBMP2/4* and *HyBMP5/8b* between iCRT14-treated regenerates and controls (Fig. 4F, G).

**Figure 4:**
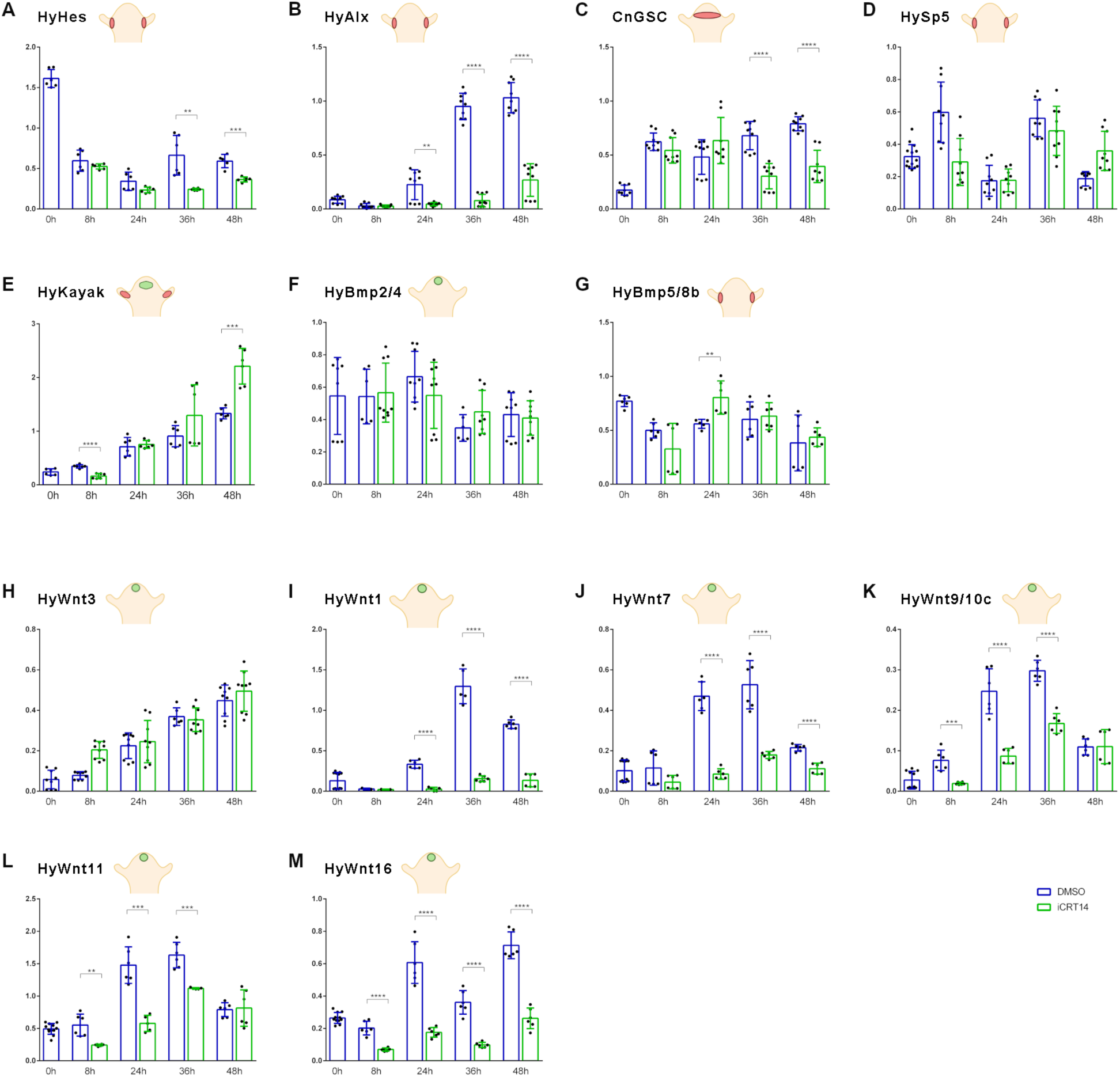
Gene expression dynamics of selected genes in iCRT14 inhibited regenerates. RT-qPCR measurements quantifying gene expression of **(A)** *HyHes* **(B)** *HyAlx* **(C)** *Goosecoid* **(D)** *Sp5* **(E)** *HyKayak* in epithelial cells, **(F)** *BMP2/4* **(G)** *BMP5/8b* **(H-M)** *Wnt3, Wnt1, 7, 9/10c, 11 and 16* during 48 hrs of *Hydra* head regeneration in iCRT14 (green) or DMSO for control (blue). *Hydra* cartoons indicate gene expression patterns according to published in situ hybridisation data and single-cell sequencing atlas (Broun, Sokol et al. 1999, Hobmayer, Rentzsch et al. 2000, Smith, Gee et al. 2000, Reinhardt, Broun et al. 2004, Lengfeld, Watanabe et al. 2009, Münder, Käsbauer et al. 2010, Watanabe, Schmidt et al. 2014, Siebert, Farrell et al. 2019, Vogg, Beccari et al. 2019); Relative normalized expression was related to the housekeeping genes *GAPDH*, *RPL13*, *EF1alpha* and *PPIB*. Regeneration time is shown on x-axes; t=0 refers to animals immediately after the head was removed. Data is shown as mean ± SEM, *: p=0.05, **: p=0.01, ***: p=0.001, ****: p=0.0001.

Confirming FISH-images shown in Fig. 1D, *HyWnt3* was not inhibited by iCRT14 during head regeneration, it even appeared slightly up-regulated at the 8 hrs time point (Fig. 4H). In contrast to *HyWnt3*, expression of all other canonical *HyWnt* genes was inhibited by iCRT14 during head regeneration. *HyWnt1, 7* and *16* were inhibited throughout the whole regeneration period (Fig. 4I, J, M). *HyWnt9/10c* and *HyWnt11* were blocked up to 36 hrs, but their expression levels returned to control values at 48 hrs (Fig. 4K, L).

In summary, RT-qPCR analyzes show that β-catenin transcriptional activity is not required for expression of *HyWnt3* during head regeneration. However, it is involved in up-regulating the canonical *Wnt*-genes *HyWnt1, 7, 9/10, 11 and 16*. Moreover, *HyAlx* expression strongly depends on β-catenin activity. Expression of *HyHes* and *CnGsc* both seem strengthened by ββ-catenin during later regeneration stages, when β-catenin also seems to inhibit *Hykayak* expression. These effects on gene expression may be due to the failure of tentacle development in iCRT14 treated animals. In contrast, *BMP2/4, BMP 5/8* and *Sp5* do not appear to be regulated by β-catenin during head regeneration.

From these analyzes we conclude (1) Notch-signaling is responsible for sustained expression of *HyWnt3* and all canonical *HyWnt* genes during head regeneration. In addition, it is required for the expression of *BMP2/4* (Broun et al., 1999) and the suggested *Hydra* organizer gene *CnGsc,* supporting our previous experiments where DAPT-treated regenerating head tissue did not develop organizer activity (Münder et al., 2013). (2) Notch activity is required for inhibiting *HyKayak* and *HyBMP5/8b* gene expression during regeneration, which coincides with DAPT causing downregulation of the transcriptional repressor and Notch-target gene *HyHes*. (3) β-catenin transcriptional activity is not necessary to express *HyWnt3,* acquire organizer activity and form a new hypostome after head removal. However, β-catenin-dependent transcription is indispensable to express *HyAlx* and form tentacles.

## 4. HyKayak

*HyWnt3*, albeit inhibited by DAPT specifically during head regeneration, had so far not been indicated as a potential target for Notch-mediated gene activation in *Hydra* (Moneer et al., 2021; Münder et al., 2013). By analyzing the *HyWnt3*-promoter region, Nakamura et al found proximal elements similar to *Drosophila* Su(H)- and RBPJ-sites (−155 to −143, (Nakamura et al., 2011)). Notch could therefore directly activate *Wnt3*-expression. However, several repressor genes are Notch-regulated (Moneer et al., 2021). We thus considered the possibility, that a repressor of *HyWnt3* could be inhibited by Notch-signaling especially at the tip of regenerating heads.

According to our previous report, the *Hydra* fos-homolog *HyKayak* (t5966aep) was up-regulated after Notch-inhibition with DAPT (Moneer et al., 2021). This suggests that *HyKayak* may serve as a potential target gene for Notch-regulated repressors including *HyHes* and *CnGsc,* and in this way *HyKayak* may be inhibited when these repressors are activated by Notch-signaling.

Analysis of domain structure and sequence comparison of *HyKayak* with *fos*- and *jun*-sequences from *Aurelia aurita*, *Stylophora pistillata*, *Caenorhabditis, Drosophila*, mouse and human revealed a strong conservation of the bZiP-domain (basic leucine zipper domain), which is responsible for DNA-binding and dimerisation (Fig. S1A, B). Phylogenetic analysis showed that *HyKayak* is related to *c-fos*-sequences of various species including *Hydra* (Fig. S1C). *HyKayak* is expressed in ectodermal cells of *Hydra* head, tentacles and body column, excluding the basal disk (Fig. S1D) (Siebert et al., 2019)). A second *fos*-gene described by Cazet et al (Cazet et al., 2021) is expressed in epithelial cells and gland cells (referred to as fos_Cazet). Additionally, we identified two transcripts encoding Jun-related proteins, HyJun_nem (t17964aep) expressed in nematoblasts 3-5, and HyJun_epi (t19405aep) expressed in all cells, with especially high levels in epithelial cells (Fig. S1D). By SDS-PAGE of *Hydra* lysates and staining with anti-*HyKayak* antibody (see Fig. S1E-a), we found that *HyKayak* protein remained in the pellet fraction and only a small percentage could be solubilized after treatment with DNase, suggesting that *HyKayak* is strongly associated with DNA and lending support to its suggested role as a DNA-binding protein (Fig. S1E-b).

Fos-proteins interact with Jun-proteins (also bZIP domain proteins) to form the transcriptional regulation complex AP-1 (activator protein 1) (Karin et al., 1997). To test such interactions for the *Hydra* proteins, we performed immunoprecipitation of HyKayak and HyJun_epi-proteins expressed in HEK293T-cells. This revealed that HyKayak did not interact with itself, but strongly interacted with HyJun_epi protein (Fig. S2). To investigate the function of HyKayak/AP-1 in *Hydra* head regeneration, we used the Fos/jun-inhibitor T5224 to block DNA-binding of Fos/Jun-complexes (Xiong et al., 2022) and analyzed gene expression and phenotypes during *Hydra* head regeneration. This revealed mild inhibition of *HyKayak* expression in contrast to strong up-regulation of *HyJun* (Fig. 5A, B). In addition, we discovered that *HyWnt3* expression was strongly up-regulated by T5224 (Fig. 5C).

**Figure 5:**
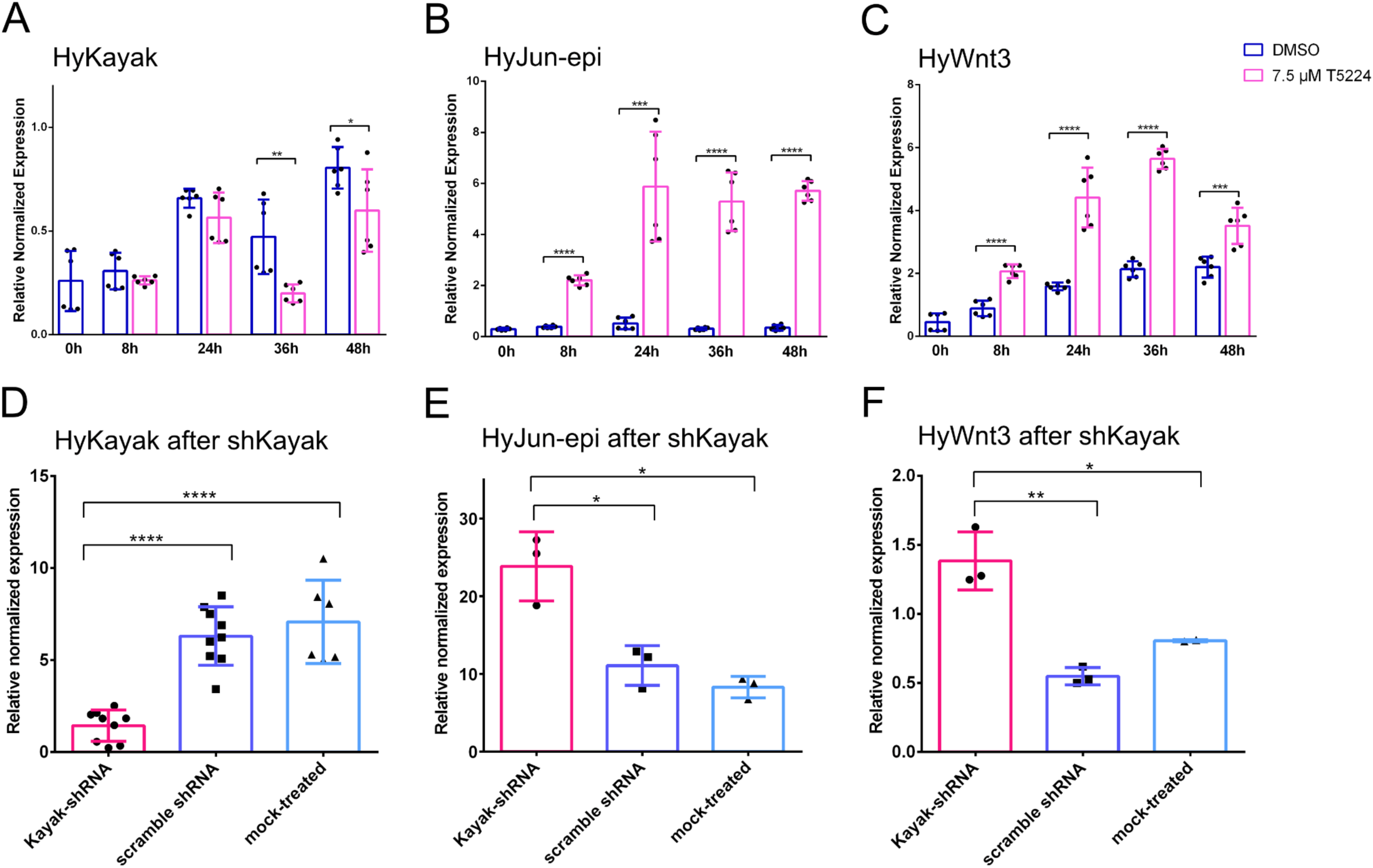
Gene expression dynamics of selected genes in *T5224* inhibited regenerates corresponds with gene expression in kayak knockdown polyps. RT-qPCR measurements quantifying expression of **(A)** *HyKayak,* **(B)** *HyJun_epi* and **(C)** *HyWnt3* for 48 hrs after head removal in HM with DMSO (control), (blue) and HM with 7.5 µM T5224 for inhibition of Fos/AP1 activity (pink). RT-qPCR measurements quantifying expression of **(D)** *HyKayak,* **(E)** *HyJun-epi* and **(F)** *HyWnt3* after knockdown *of HyKayak* by shRNA (pink), compared to control knockdown with scramble shRNA (dark blue) and mock-knockdown control (light blue). Relative normalized expression was related to the housekeeping genes GAPDH, EF1alpha and PPIB. Regeneration time is shown on x-axes; t=0 refers to animals immediately after the head was removed. Data is shown as mean ± SEM, *: p=0.05, **: p=0.01, ***: p=0.001, ****: p=0.0001.

To confirm that *HyKayak* was involved in *HyWnt3* overexpression, we knocked down *HyKayak* using shRNA directed against *HyKayak*. We achieved *HyKayak* knockdown by ca. 80% in comparison with control polyps either mock-treated or treated with scrambled control shRNA (Fig. 5D). Moreover, Kayak-knockdown led to the upregulation of *HyJun*, consistent with the effects of T5224 treatment (Fig. 5E). Importantly, knockdown of *HyKayak* induced up-regulation of *HyWnt3* (Fig. 5F). From these data we conclude that (1) *HyKayak* attenuates expression of *HyWnt3*; (2) HyKayak may work within the AP-1 complex together with Jun-epi and (3) Notch signaling may block the inhibitory activity of *HyKayak* on *HyWnt3* by activating repressor genes. With DAPT, *HyKayak* remains active and inhibits sustained expression of *HyWnt3* at later stages of head regeneration.

## 5. Regeneration of Craspedacusta polyps

Our data dissect the regeneration of *Hydra* heads into two processes, formation of the hypostome and head, and formation of tentacles. For hypostome formation *HyWnt3* is needed but β-catenin transcriptional activity is dispensable. Notch-signaling then appears to be responsible to “organize” these two morphogenetic processes. In order to test this hypothesis, we asked, how inhibition of Notch-signaling might affect regeneration of polyps with a simpler, one-component, head. We used polyps of the fresh water hydrozoan *Craspedacusta sowerbii.* They have a mouth opening that is surrounded by epithelial cells carrying nematocytes, but they don’t possess tentacles (Ramos et al., 2017).

*Craspedacusta* polyps are shown in Figure 6A. They often occur as mini-colonies with one foot carrying two polyps. Actin fibres are running along the polyp’s body column and form a ring where the two polyps separate just above the foot. Actin cushions carrying nematocysts are visible and indicate the positions of capsules along the body column and in a ring surrounding the mouth opening (Fig. 6A, B). Additional capsule staining with DAPI (Szczepanek et al., 2002) very clearly reveals the pattern of nematocysts in the head (Fig. 6B). When we removed the heads of the polyps, most of them fully regenerated within 96 hrs (Fig. 6B). Some retracted into a podocyst (the ‘dauerstadium’) (Fig. 6D). Polyps treated with DMSO or DAPT also completed head regeneration after 96 hrs (Fig. 6C). Quantification of *Craspedacusta* development after head removal revealed that similar numbers of proper head regeneration and podocyst formation occurred (Fig. 6D). This indicated that Notch-signaling was not required for head regeneration in *Craspedacusta-*polyps.

**Figure 6:**
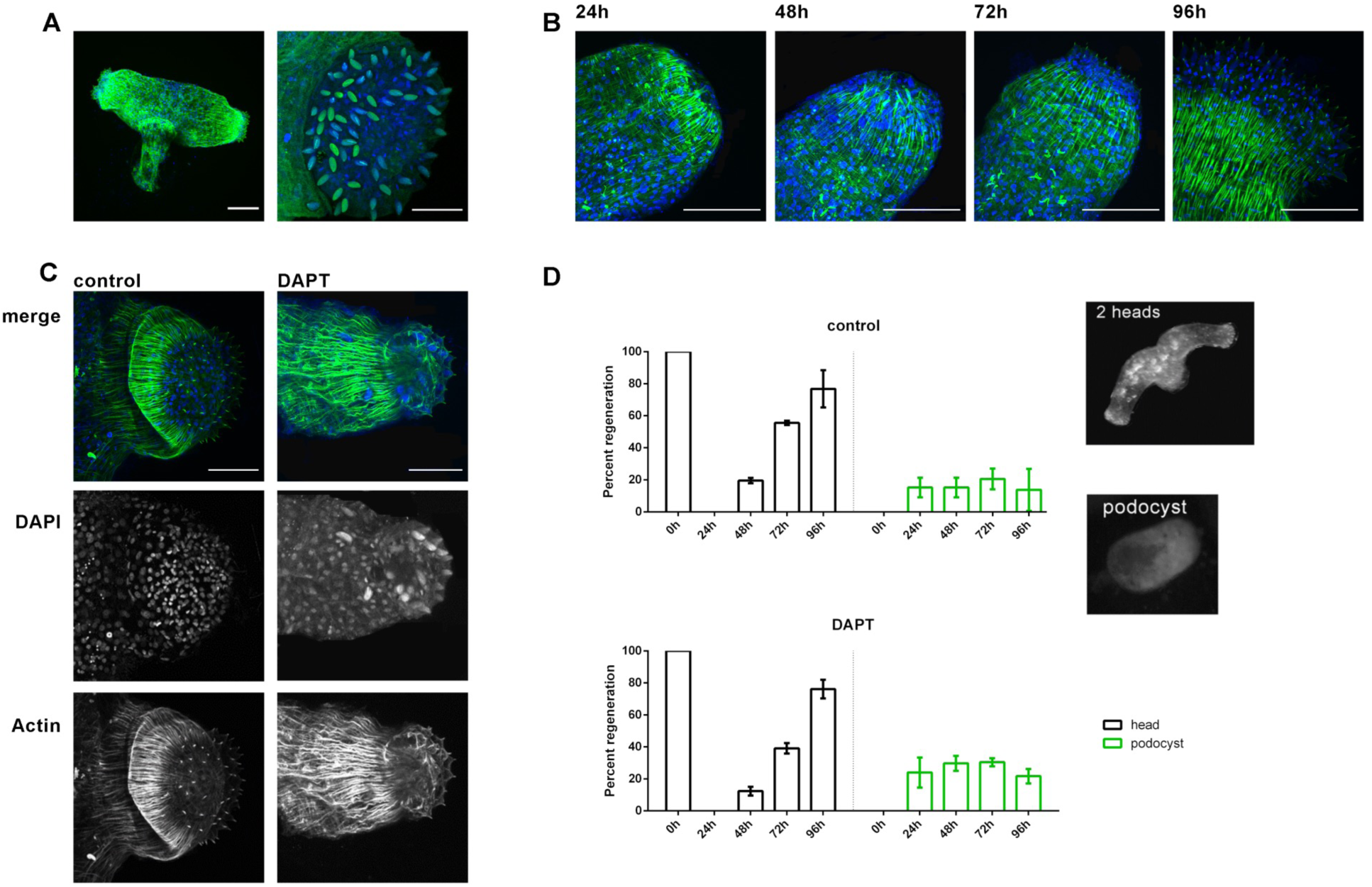
Head regeneration of *Craspedacusta* polyps. **(A)** *Craspedacusta* colony with two animals sharing one foot, Scale bar: 100 µm. Magnification of polyp head with nematocytes surrounding the hypostome, Scale bar: 50 µm. Nematocyte capsules stained with DAPI (green). **(B)** Regeneration of *Craspedacusta* polyps at 24, 48, 72 and 96 hrs after head removal with reappearing nematocytes and actin fibres. Scale bar: 50 µm. **(C)** Regenerated heads 96 hrs after head removal in HM with DMSO (control) or DAPT as indicated. Scale bar: 50 µm. Actin fibres are shown in green after staining with FITC-phalloidine, nuclei are shown in blue after staining with DAPI. **(D)** Percentage of *Craspedacusta* polyps showing normal regeneration with 1, 2 or 3 heads (black) or ‘podocyst’ form (green) at indicated time points after head removal. Representative images of *Craspedacusta* polyps showing a colony with 2 heads (upper panel) and a podocyst (lower panel).

In order to confirm that DAPT was taken up by the polyps even in the absence of a visible regeneration phenotype, we investigated the effect of the drug under regeneration conditions on expression of some possible Notch-target genes. We choose homologs of *HyAlx* and *HySp5*, both genes had been identified as Notch-target genes in *Hydra* (Moneer et al., 2021), and a homolog of *NOWA,* a gene encoding a protein of the outer nematocyte capsule wall (Figs. S5-S7). In *Hydra*, *NOWA* is down-regulated by DAPT because of the defect in nematocyte differentiation, which occurs when Notch-signaling is blocked (Käsbauer et al., 2007; Moneer et al., 2021). The results are shown in Fig. 7. DAPT inhibits the expression of *CsAlx* and of *CsSp5* during head regeneration. It also inhibits the expression of *CsNOWA*. This effect of DAPT on the expression of homologs of suggested *Hydra* Notch-target genes confirms that the drug must have entered the cells in *Craspedacusta* polyps.

**Figure 7:**
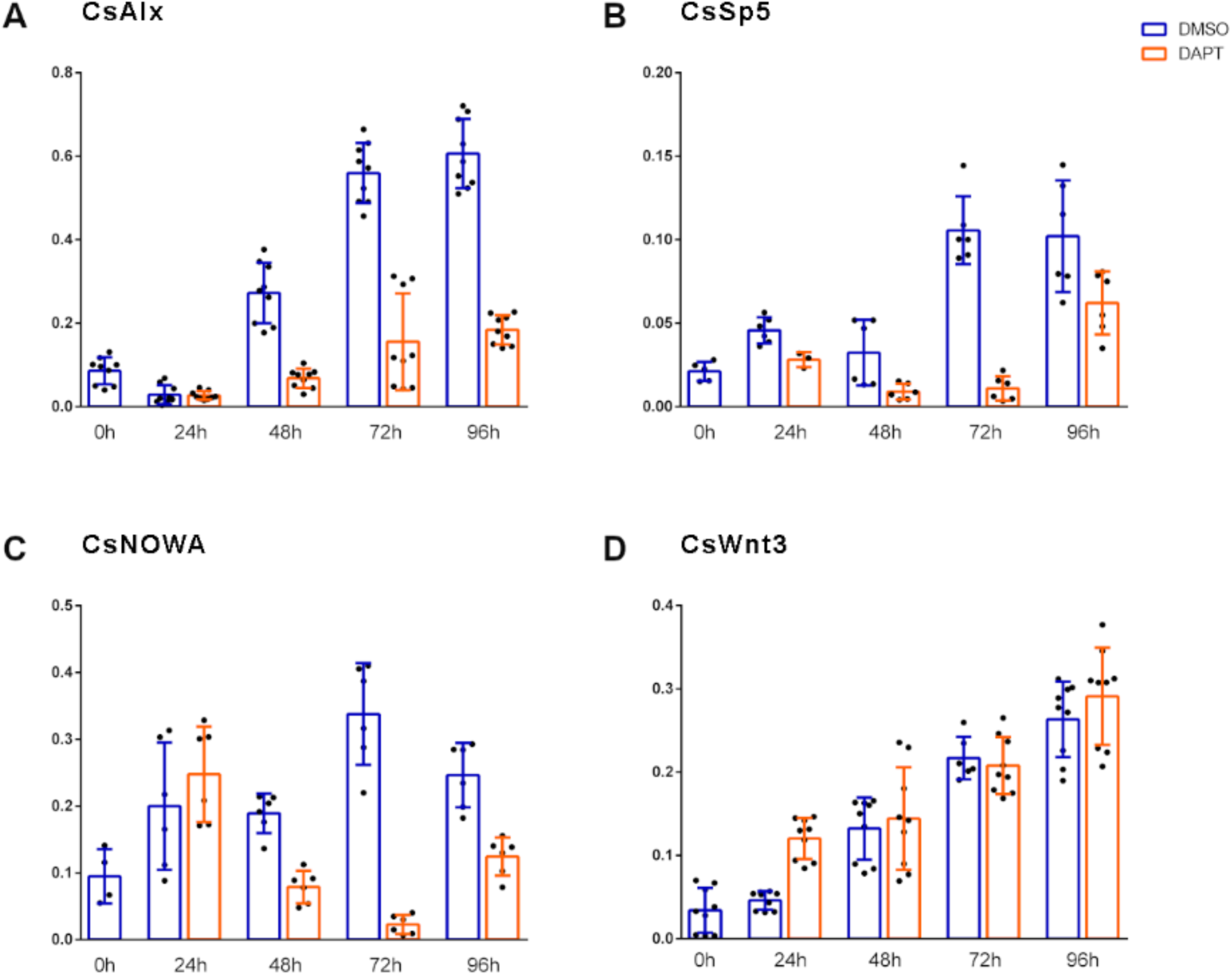
Gene expression dynamics of selected genes in DAPT inhibited regenerates of Craspedacusta. RT-qPCR measurements quantifying expression of *Craspedacusta* genes **(A)** *CsAlx,* **(B)** *CsSp5,* **(C)** *CsNOWA,* **(D)** *CsWnt3* during 96 hrs of head regeneration in DAPT (orange) or DMSO for control (blue). Relative normalized expression related to the housekeeping genes *GAPDH*, *actin* and *PPIB*. Regeneration time is shown on x-axes; t=0 refers to animals immediately after the head was removed. Data is shown as mean ± SEM, *: p=0.05, **: p=0.01, ***: p=0.001, ****: p=0.0001.

Finally, we investigated the expression of the *Craspedacusta Wnt3-*gene (Fig. 7) and its response to DAPT treatment during head regeneration. We observed a low expression level of *CsWnt3* after head removal (t=0), which dramatically increased as the head regenerated, suggesting that *Wnt3* is expressed in the head of *Craspedacusta* polyps like in the head of other cnidarians, including *Hydra*, *Hydractinia* and *Nematostella* (Hobmayer et al., 2000; Kusserow et al., 2005; Plickert et al., 2006). Consistent with its lack of effect on head regeneration, DAPT also did not inhibit *CsWnt3* expression during this process in *Craspedacusta*. This is opposite to the situation in *Hydra.* If *CsWnt3* would be involved in *Craspedacusta* head regeneration, this could explain the failure of DAPT to interfere with this process.

## Discussion

Head regeneration in *Hydra* can be divided into two processes, re-formation of a hypostome-body column-axis and re-formation of tentacles. We show here that tentacle formation requires β-catenin transcriptional activity, but hypostome regeneration does not. Conversely, hypostome regeneration requires Notch-signaling, whereas tentacle tissue does not. By RT-qPCR gene expression analysis we investigated the expression dynamics of selected genes in response to inhibition of β-catenin transcriptional activity, or of Notch-signaling over a regeneration time of 48 hrs in polyps after heads had been removed at an apical position, just underneath the tentacles.

The results of these gene expression analyses are schematically displayed in Table 1. We distinguish two phases of regeneration, the first 8 hrs and the time thereafter. With the exception of the direct Notch-target gene *HyHes* (Moneer et al., 2021; Münder et al., 2013) expression of our selected genes is not affected by DAPT 8 hrs after head removal. This time is allocated to wound healing, and this process appears independent of Notch-signals (Cazet et al. 2021). However, over the following time course expression levels of *HyWnt-*1, 3, 7, 9/10, 11 and 16, all implied in canonical Wnt-signaling, declined to almost zero in DAPT treated polyps. In addition, the potential “organizer” gene *CnGsc* was inhibited with DAPT corresponding to the observation that Notch-treated regenerates do not acquire organizer activity. Sp5, which was suggested to be part of an inhibition loop for *HyWnt3-*β-catenin (Vogg et al., 2019) and a direct Notch-target gene (Moneer et al., 2021) was also blocked by DAPT during head regeneration. *HyAlx*, which has repeatedly been shown to induce differentiation of tentacle tissue (Broun and Bode, 2002; Broun et al., 2005; Gee et al., 2010; Münder et al., 2013; Smith et al., 2000) was only slightly affected by DAPT, corresponding to the detection of irregular tentacles in some regenerates (Münder et al., 2013). However, the lack of organizer activity in such regenerates may be responsible for their failure to produce correct tentacle patterns. We also observed that the expression of *HyBMP2/4* is strongly dependent on Notch-signaling. Together these results suggest that *Hydra* head regeneration requires canonical *Wnt*- and *BMP2/4*-signaling to produce organizer and hypostome, both of which depend on the presence of Notch-signaling. In contrast, *HyBMP5/8b* and *HyKayak* were up-regulated by DAPT, suggesting that Notch was required to inhibit these genes.

**Table 1:**
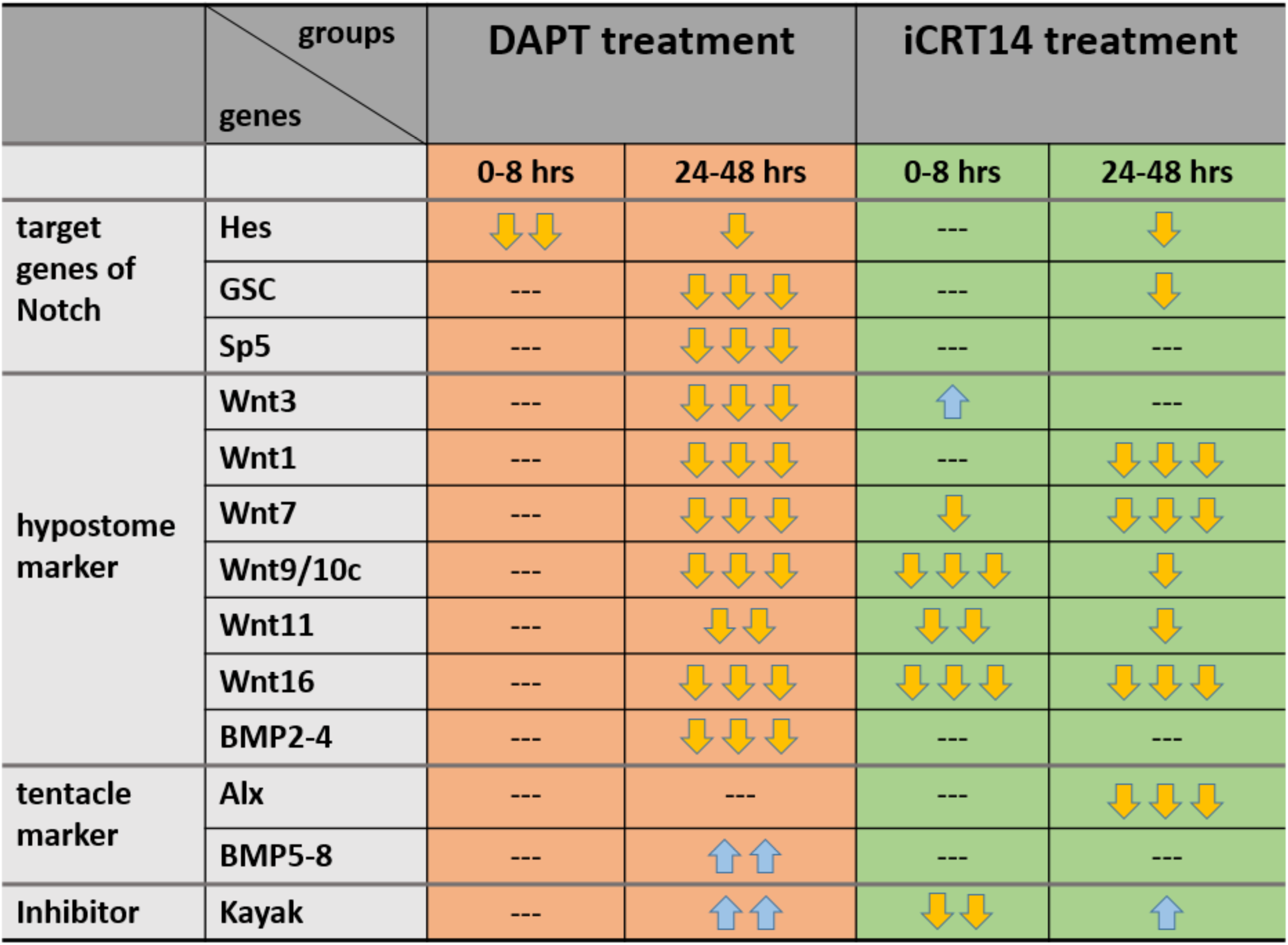
Summary of changes in gene expression during *Hydra* head regeneration in medium with Notch- and β-catenin inhibitors. RT-qPCR results indicating effect of inhibition of Notch-signaling by DAPT (orange) and inhibition of β-catenin transcriptional activity by iCRT14 (green) on expression of *HyHes, CnGsc, HySp5, HyWnt3, 1, 7, 9/10c, 11, 16, HyBMP2/4, HyBMP5/8, HyAlx* and *HyKayak*. Gene expression is classified within the initial first 8 hrs and between 24-48 hrs of regeneration; up-regulation is indicated by blue arrows, downregulation by yellow arrows. Numbers of arrows refers to the strength of the effect; dotted lines mean no effect.

We also found that tentacle tissue formation, especially the expression of *HyAlx* in apical regenerates was completely blocked with iCRT14. On the contrary, it is known that increasing nuclear β-catenin (and thus its transcriptional activity) by alsterpaullone induces formation of ectopic tentacles, but not hypostomes or even complete heads (Broun et al., 2005). Therefore, the phenotype observed with iCRT14 is obviously caused by a lack of tentacle activation, while ectopic activation of β-catenin induces tentacle formation through activation of *HyAlx*.

Most intriguingly, induction of *HyWnt3* expression in apical regenerates was not blocked in the absence of β-catenin transcriptional activity, indicating that *HyWnt3* is not up-regulated via β-catenin dependent autoactivation after head removal, as had been suggested to occur in un-disturbed polyps (Nakamura et al., 2011). In contrast to *HyWnt3*, all other canonical *Wnt*-genes were downregulated by iCRT14, at least to some extent, indicating that they were β-catenin-dependent. In the presence of iCRT14 *HyWnt3* must perform its function during head regeneration by signaling through a β-catenin independent pathway. Remarkably, iCRT-treated tissue regenerated perfect hypostomes with the normal *HyWnt3* expression pattern.

The effect of iCRT14 had been analyzed in previous studies (Cazet et al., 2021; Gufler et al., 2018; Tursch et al., 2022). All showed β-catenin-dependency for down-regulation of head specific genes in foot regenerates at time points up to 12 hrs after head removal, including *HyWnt3*. They also stated a failure of head regeneration in the presence of iCRT14 but, in accordance with our study, did not reveal that *HyWnt3* expression at future heads depended on β-catenin. None of these studies analyzed the regeneration of tentacles and hypostomes separately and they did not report whether the regeneration of hypostomes 48 hrs after head removal occurred normally upon iCRT14 treatment.

While the tissue left after head removal has the capacity to form both, tentacles and hypostome/head, final patterning of the new head involves emergence of hypostome and tentacle structures at distinct locations. A model proposing two independent patterning systems, each comprising an activator and an inhibitor for head and tentacle formation, had been introduced before, when *HyAlx* was discovered (Smith et al., 2000). After cutting off the head at apical positions, *HyAlx* first appeared at the tip. This was explained with high tentacle activation potential in this region, leading to a fast establishment of the tentacle system with *HyAlx* expression and tentacle markers (like *HMMP*) covering the whole regenerating tip. Tentacle activation is then inhibited by a tentacle inhibitor. Head activation takes over and expression of canonical *Wnt*-genes becomes stronger. *HyAlx* shifts to the emerging tentacle region and finally appears in rings from which tentacles emerge (see Fig. 8).

**Figure 8:**
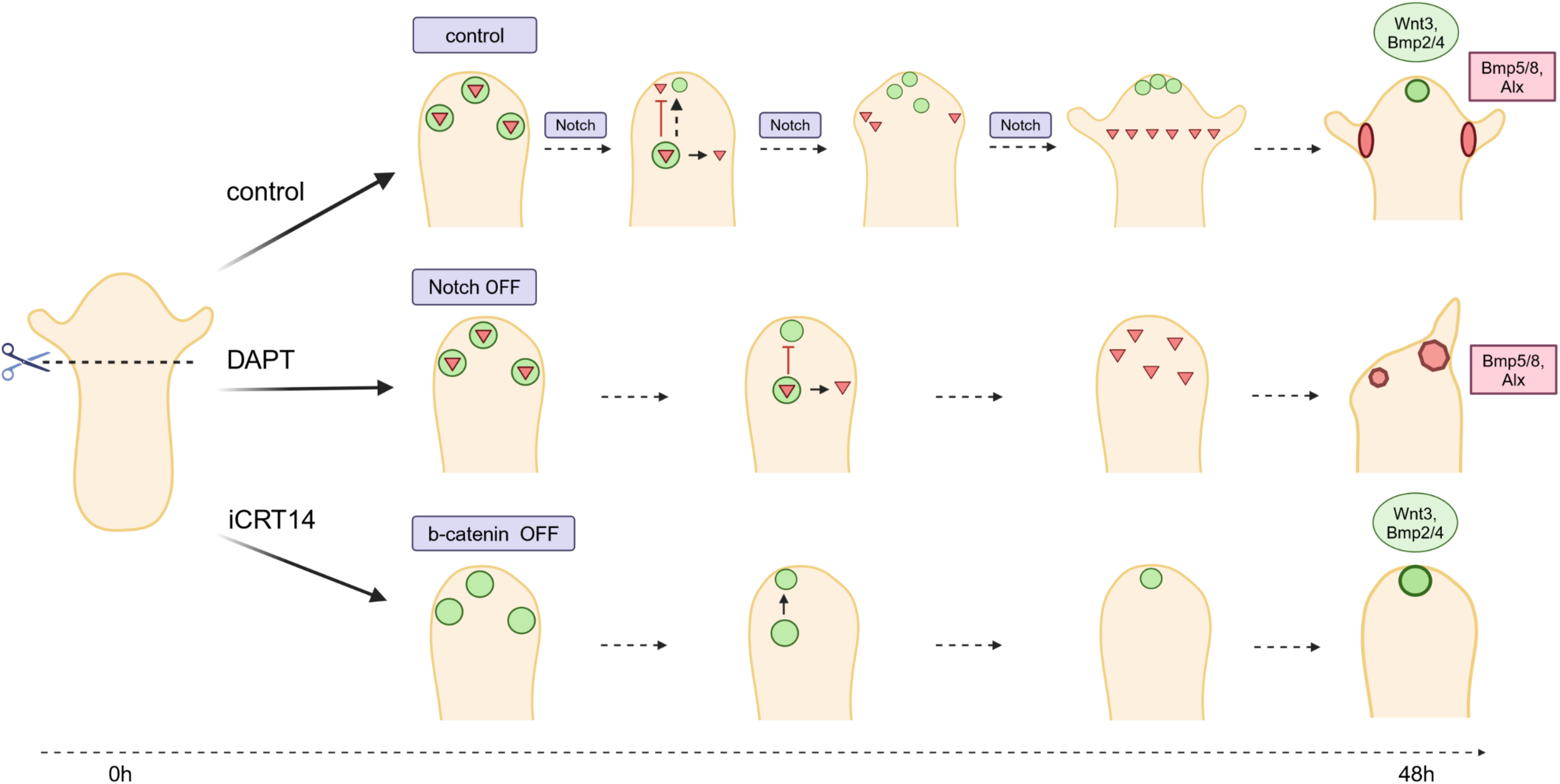
Model for Notch-function in *Hydra* head regeneration in comparison with β-catenin. Schematic representation of DAPT- and iCRT14 effects on *Hydra* head regeneration. Hypostome is labelled green; tentacle boundary red (red). Animals treated with DAPT regenerate tentacle boundary gene expression in irregular patterns and show irregular tentacle morphology. Animals treated with iCRT14 regenerate regular expression pattern of hypostomal genes (HyWnt3) and show normal hypostome morphology. They do not regenerate any HyAlx-expression and do not show tentacles. Model for course of *Hydra* head regeneration in the presence of Notch-signaling (control, upper panel), absence of Notch-signaling (middle panel) and absence of Wnt-signaling (lower panel). After head removal the potential to re-form a head (green circles) and tentacles (red triangles) arises in the regenerating tip of the polyp. Notch-signaling then mediates inhibition of the tentacle fate in the upper part of the regenerate by repressing HyBMP5/8 and allowing expression of HyBMP2/4 and HyWnt3 (hypothetically by repressing inhibitors of these genes as indicated by dotted line). This allows re-establishment of the hypostome and organizer tissue, while confining tentacle development to the lower part of the regenerate (control). With inhibition of Notch (middle panel) tentacle fate is not inhibited in the tip of the regenerate preventing expression of hypostomal genes (HyWnt3 and HyBMP2/4) and allowing tentacle tissue development in the whole regenerating tip. However, as an organizer does not develop, this tissue cannot be patterned properly (red triangles). Without transcriptional activity of β-catenin hypostomal genes (Wnt3 and HyBMP2/4) are expressed, while tentacle tissue is not induced (no HyAlx). Created with Biorender.

In contrast, budding starts with head activation being established and *HyAlx* is expressed later, always excluding the apical part of the bud. This was attributed to higher head activation potential in the budding region in comparison with tentacle activation activity. Moreover, older regeneration experiments had revealed that apical and basal regenerates differed in the order of appearance of head and tentacle tissue. Tentacle tissue appeared first in apical regenerates and later in basal ones (Technau and Holstein, 1995).

Here we have only considered apical regenerates where the heads of the polyps were cut off just underneath the tentacles. We suggest that Notch-signaling fulfills a role in tentacle inhibition in this case. Without this inhibition, head activation with expression of all canonical *Wnt*-genes does not occur. However, Notch also affects head regeneration at basal cuts, as we have recently shown by analyzing transgenic *Hydra* with inhibited Notch-function. Here a substantial part of the animals regenerated two heads (Pan et al., 2024). This again confirms the idea that head and tentacle formation use two independent patterning systems and Notch is required to mediate between or “harmonize” them. When the tentacle system is activated first, Notch inhibits it to allow emergence of the head system. When the head system emerges first, Notch blocks it to prevent the formation of multiple heads.

How does tentacle inhibition work? It is well established that Notch activates transcriptional repressors, including *HyHes* genes and thereby suppresses specific cell fates in signal receiving cells, whereby allowing those fates in signal sending-cells (Bray, 2006). Our data suggest, that DAPT inhibited the expression of two established transcriptional repressor genes, *HyHes* and *CnGsc*. This poses the question for targets of these repressors, which should be up-regulated when Notch-signaling is inhibited. We observed this behavior for *BMP5/8b* and *HyKayak.* On the basis of the published *BMP5/8b* expression patterns (Reinhardt et al., 2004), this gene is probably part of the tentacle patterning system.

*HyKayak* encodes a homolog of Fos-proteins, which are components of the AP1 transcriptional complex, as we show by sequence comparison and phylogenetic analysis of the bZIP-domain. Moreover, HyKayak interacted with HyJun, but not with itself, similar to the behavior of human c-Fos, which does not form homodimers but instead heterodimerizes with Jun-proteins (Kouzarides and Ziff, 1988). Fos is suggested to be a negative regulator of its own promoter (Sassone-Corsi et al., 1988), and Fos can function as a repressor on cellular immediate-early genes, like Egr genes (Gius et al., 1990). Both repressions are mediated by the C-terminus of Fos and are independent of Jun (Gius et al., 1990; Ofir et al., 1990). However, the C-terminus of Fos is not required for the repression of cardiac transcription and muscle creatine kinase (MCK) enhancer (Lassar et al., 1989; Li et al., 1992; McBride et al., 1993). Our hypothesis that *HyKayak* could repress the *HyWnt3* gene was confirmed by shRNA-mediated *HyKayak*-knockdown, which resulted in the up-regulation of *HyWnt3* expression. In addition, *HyJun-epi* was also up-regulated. This is in accordance with previously published observations in human prostate cell lines where *fos*-loss-of-function has resulted in up-regulation of *jun*-expression (Riedel et al., 2021). Moreover, experiments with pharmacological inhibition of the AP1-complex with T5224 during head regeneration revealed that *HyWnt3* and *HyJun-epi* were strongly up-regulated. We therefore suggest that the *Hydra fos*-homolog *HyKayak* inhibits *HyWnt3* expression and can be a target for a Notch-induced transcriptional repressor (such as *HyHes*) in the regenerating *Hydra* head. Nevertheless, we were not able to rescue the DAPT-phenotype by inhibiting HyKayak, neither by using the inhibitor nor by shRNA-treatment, probably due to the strength of the DAPT effect. Therefore, we cannot exclude the possibility that Notch activates *HyWnt3* directly or that it represses unidentified Wnt-inhibitors through *HyHes* or *CnGsc*.

Different bZiP transcriptional factors (TFs) may have different effects on the expression of Wnt genes, and these effects are context-dependent. In previous research, Cazet et al. identified another *Hydra fos* gene (referred to as fos_cazet) and bZiP TF binding sites in the putative regulatory sequences of *HyWnt3* and *HyWnt9/10c*. They showed that bZiP TF-genes, including *Jun* and *fos*, were transiently upregulated 3 hrs after amputation, therefore they hypothesized that bZiP TFs could induce TCF-independent upregulation of *HyWnt3* during the early generic wound response (Cazet et al., 2021). However, in our study *HyKayak* expression continuously increased throughout the entire head regeneration process (Fig. 3E and 4E) including the morphogenesis stages (24-48 hrs post-amputation). Another study reported that inhibition of the JNK pathway (which disrupts the formation of the AP-1 complex) resulted in upregulation of *HyWnt3* expression in both, head and foot regenerates (Tursch et al., 2022). This result might support our hypothesis, but it only included the first 6 hours after amputation, similar to Cazet’s research. Therefore, it appears that *HyKayak* and fos_Cazet may have opposing roles in the regulation of *Wnt*-gene expression and are possibly activated by different signaling pathways depending on the stages of regeneration.

Notch activity is dependent on the regeneration time. At early time points it is apparently not required, but between 8 and 48 hrs after head removal loss of Notch activity severely impairs the regeneration process (Fig. 3, Fig. 8A, B). In addition, the gene expression dynamics for many of the analyzed genes appears in wave-like patterns in some experiments (see Figs S3 and S4). As we have only four time-points measured, we cannot draw strong conclusions from these observations, except that some of the deviations in our data points (e.g. 48 hrs *HyHes*) might be caused by oscillations. It is tempting to propose that the dynamic development of gene expression patterns over the time course of regeneration hints at a wave like expression of Notch-target genes (e.g. *HyHes*). *Hes*-genes have been implicated in mediating waves of gene expression, e.g. during segmentation and as part of the circadian clock (Kageyama et al., 2007). This property is due to the capability of Hes-proteins to inhibit their own promoter. Future models for head regeneration in *Hydra* should consider the function of Notch to inhibit either module of the regeneration process and the potential for the Notch/Hes system to cause waves of gene expression. Such waves intuitively seem necessary to change the gene expression patterns underlying morphogenesis during the time course of head regeneration.

Is Notch part of the organizer? The organizer is defined as a piece of tissue with inductive and structuring capacity. Notch is expressed in all cells of *Hydra* polyps (Prexl et al., 2011) and overexpression of NICD does not induce second axes all over the *Hydra* body column (Pan et al., 2024), as seen with overexpression of stabilized β-catenin (Gee et al., 2010). Moreover, Notch functions differently during regeneration after apical and basal cuts. Phenotypically during head regeneration in DAPT, we clearly recognize a missing inhibition of tentacle tissue after apical cuts and diminished inhibition of head induction after basal cuts (Pan et al., 2024).

We would thus rather suggest that the organizer activity of *Hydra* tissue utilizes Notch-signaling as a mediator of inhibition. As our study of transgenic NICD overexpressing and knockdown polyps had suggested, the localization of Notch signaling cells depends on relative concentrations of Notch- and Notch-ligand proteins, which are established by gradients of signaling molecules that define the *Hydra* body axis (Pan et al., 2024; Sprinzak et al., 2010). This is in very good agreement with the “reaction-diffusion-model” provided by Alfred Gierer and Hans Meinhardt (Gierer and Meinhardt, 1972; Meinhardt and Gierer, 1974) suggesting a gradient of positional values across the *Hydra* body column. This gradient may determine the activities of two activation/inhibition systems, one for tentacles and one for the head. When the polyps regenerate new heads, Notch could provide inhibition for either system, depending on the position of the cut.

Head regeneration also occurs in the colonial sea water hydrozoan *Hydractinia*. Colonies consist of stolons covering the substrate and connecting polyps, including feeding polyps, which have hypostomes and tentacles, and are capable of head regeneration, similar to *Hydra* polyps. *Wnt3* is expressed at the tip of the head and by RNAi mediated knockdown it was shown that this gene is required for head regeneration (Duffy et al., 2010). In the presence of DAPT, proper head regeneration did not occur, similar to *Hydra*. However, regeneration of the nerve ring around the hypostome was observed, indicating the possibility that hypostomes had been regenerated. Unfortunately, this study did not include gene expression data and therefore it is not clear whether *Wnt3* expression was affected or not (Gahan et al., 2017).

An interesting question was whether regeneration of cnidarian body parts, which are only composed of one module, also requires Notch-signaling. This is certainly true for the *Hydra* foot, which regenerates fine in the presence of DAPT (Käsbauer et al., 2007). Moreover, we tested head regeneration in *Craspedacusta* polyps, which do not have tentacles, and showed that DAPT does not affect this regeneration process. This corroborates our idea that Notch is required for regeneration in cnidarians, when this process involves two pattern forming processes to produce two independent structures, which are controlled by different signaling modules. This would be the case for the *Hydra* and for the *Hydractinia* heads, but not for *Craspedacusta*.

Future studies on expression patterns of the genes that control formation of the *Hydra* head, including *Sp5* and *Alx* in *Craspedacusta* could provide new insights into the evolution of cnidarian body patterns. *Sp5* and *Alx* appear to be conserved targets of Notch-signaling in the two cnidarians we have investigated. *Wnt3*, while being inhibited by Notch-inhibition in *Hydra* head regenerates, is not a general target of Notch signaling. It was not affected by DAPT in our comparative transcriptome analysis (Moneer et al., 2021) on uncut *Hydra* polyps, and it was also not affected by DAPT in regenerating heads of *Craspedacusta*.

## Acknowledgements

We want to express our gratitude to the funding agencies for supporting this work. A.B. and L.S. are funded by the German Research foundation (DFG) projects BO1748-12 and 17, Q.P. is funded by a CSC-grant (Chinese Scholarship Council), L.S. is funded by *Evangelisches Studienwerk Villigst e.V.* We thank Dr. Stefan Krebs, LMU Munich for sequencing of *Craspedacusta* cDNA and Dr. Sergio Vargas, LMU Munich for bioinformatic sequence analysis. The drawings in this document were produced using Affinity Designer 2 (Version 2.3.0) by Q.P.

## Data availability

All data presented in the main manuscript and supplementary files will be provided by the corresponding author (Angelika Böttger) upon request.

## Author contribution statement

A.B. and M.J. conceived this study. M.S., M.J., L.S. Q.P., A.de la P. and J.M. carried out quantitative RT-PCR and in situ hybridisation, L.S. provided all work on Craspedacusta, Q.P provided all work on HyKayak, while H.F. produced monoclonal anti-Kayak-antibody. M. M. provided GAM-based visualisation of gene expression, M.H. provided staining and imaging of Hydra thin sections, M.S., J.M. and A.B. drafted the manuscript. All authors revised and approved the manuscript.

---

**Figure S1:**
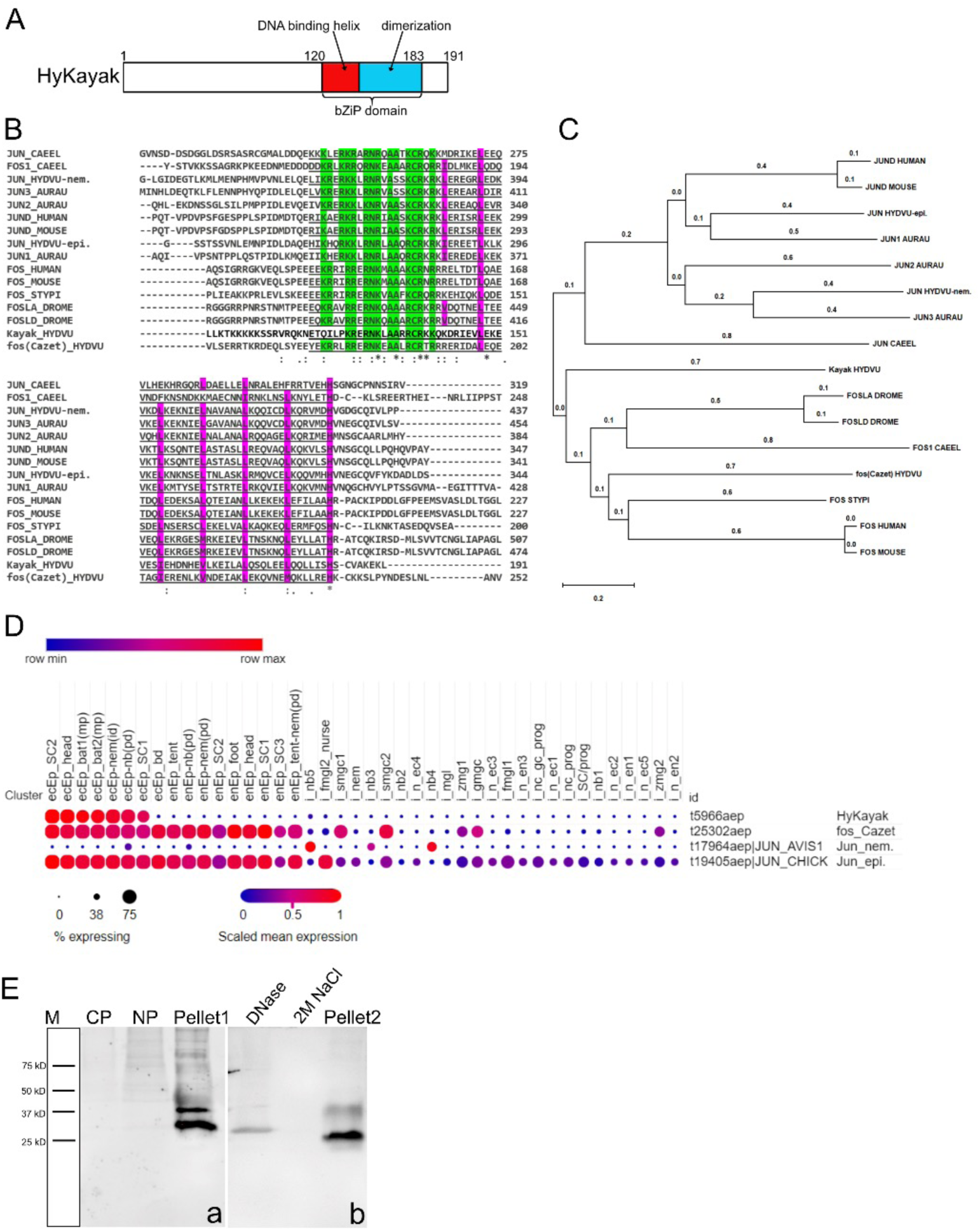
Kayak gene identification and protein domain structure. **(A)** Schematic representation of the HyKayak protein structure (191 amino acids). The bZiP domain with DNA binding and dimerization function is located between amino acids 120 to 183. **(B)** Alignment of the protein sequences of *Hydra*-Kayak, fos-Cazet, Jun-epi and Jun-nem from *Hydra Vulgaris* (HYDVU), FOS and JUN from HUMAN, MOUSE, *Caenorhabditis elegans* (CAEEL), *Aurelia aurita* (AURAU), *Stylophora pistillata (STYPI)* and *Drosophila melanogaster* (DROME); the bZIP domain is underlined, green background indicates amino acids involved in DNA-binding, violet background indicates amino acids of the dimerization interface. **(C)** Phylogenetic tree based on the alignment of 15 full-length protein sequences affiliated to the FOS and JUN families using MEGA software. Species code: *Aurelia aurita* (AURAU), *Hydra Vulgaris* (HYDVU), HUMAN, MOUSE, *Caenorhabditis elegans* (CAEEL), *Drosophila melanogaster* (DROME), *Stylophora pistillata (STYPI)***. (D)** Dot-plot of the expression patterns for the genes *HyKayak* (t5966aep), fos_Cazet (t25302aep), *HyJun_nem* (t17964aep), *HyJun-epi* (t19405aep) from the single cell portal (Siebert, Farrell et al. 2019). **(E)** Western blot was stained with anti-kayak-antibody (in-house). Lysates from *Hydra* polyps indicating cytoplasmic proteins (CP), Nuclear proteins (NP), pellets after centrifugation at 14000 g (Pellet1), supernatant after treatment of pellet with DNAse (DNase), supernatant after treatment of pellet fraction with 2 M NaCl (2 M NaCl) and pellet fraction after both treatments (pellet2).

**Figure S2:**
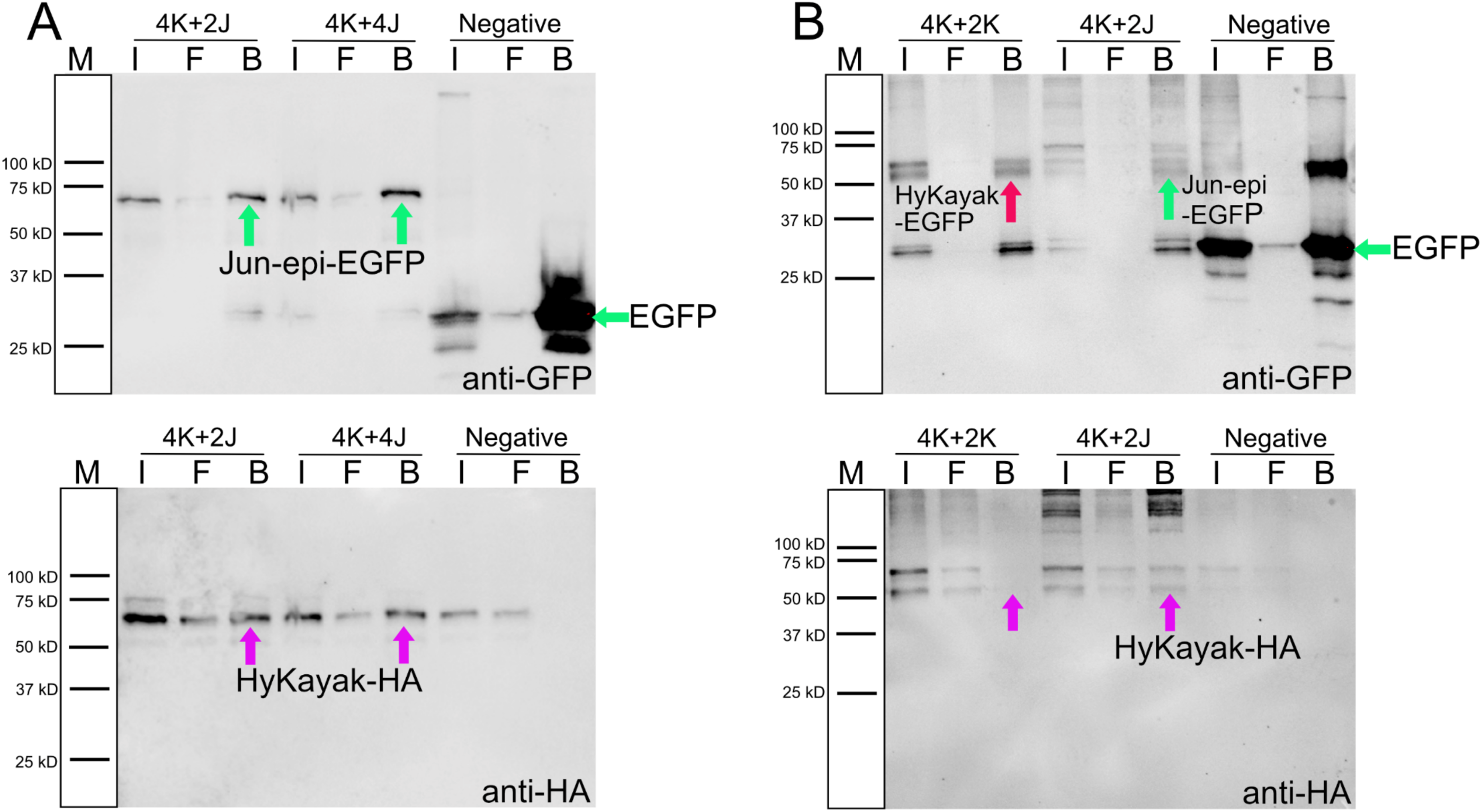
Kayak Co-immunoprecipitation. **(A)** EGFP-tagged Jun-epi was immunoprecipitated with GFP-Trap agarose beads and detected on western blot using an anti-GFP antibody. Co-precipitation of HA-tagged Kayak was detected on western blot using an anti-HA antibody in input, flow through and beads fractions. 4K+2J: 4 µM Kayak-HA plus 2 µM Jun-epi-EGFP; 4K+4J: 4 µM Kayak-HA plus 4 µM Jun-epi-EGFP. I: input; F: flow-through; B: beads. **(B)** EGFP-tagged Kayak and Jun-epi were immunoprecipitated by GFP-Trap agarose beads and detected on western blot using an anti-GFP antibody. Co-precipitation of HA-tagged Kayak was detected on western blot using an anti-HA antibody in input, flow through and beads fractions. 4K+4J: 4 µM Kayak-HA plus 4 µM Jun-epi-EGFP; 4K+2J: 4 µM Kayak-HA plus 2 µM Jun-epi-EGFP, which was used for positive control; the empty plasmid pEGFP-C1 was used for negative control.

**Figure S3:**
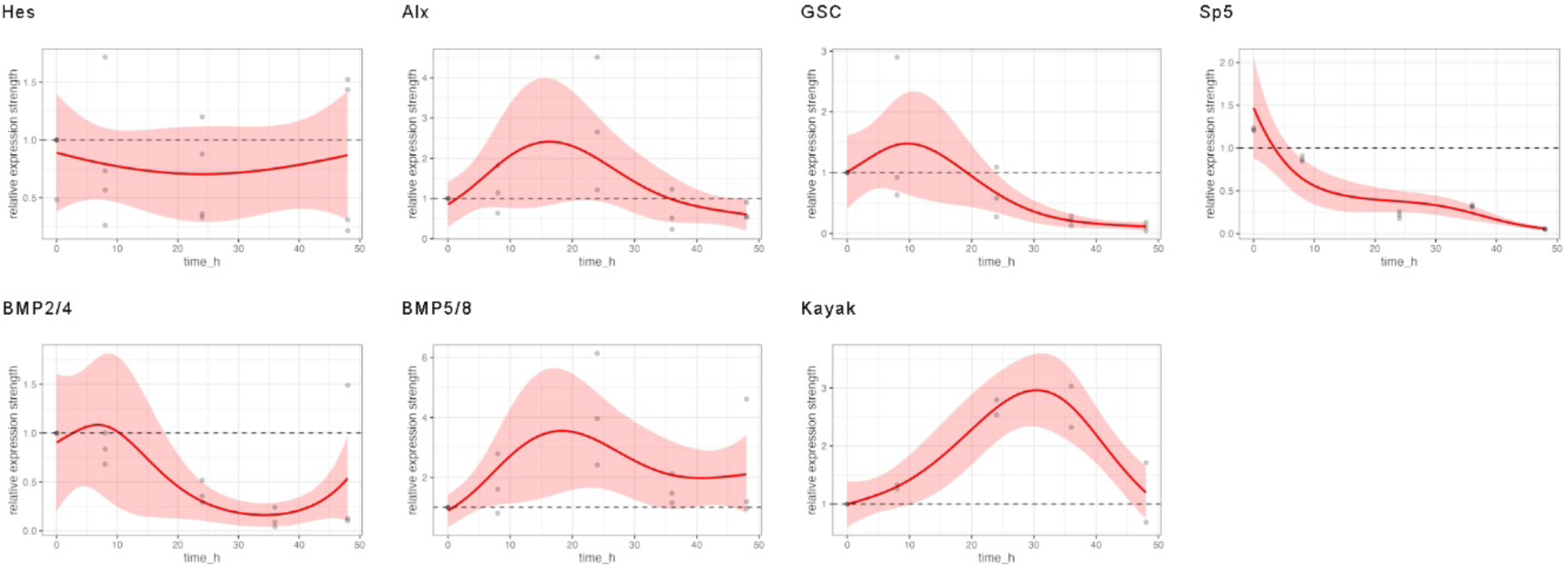
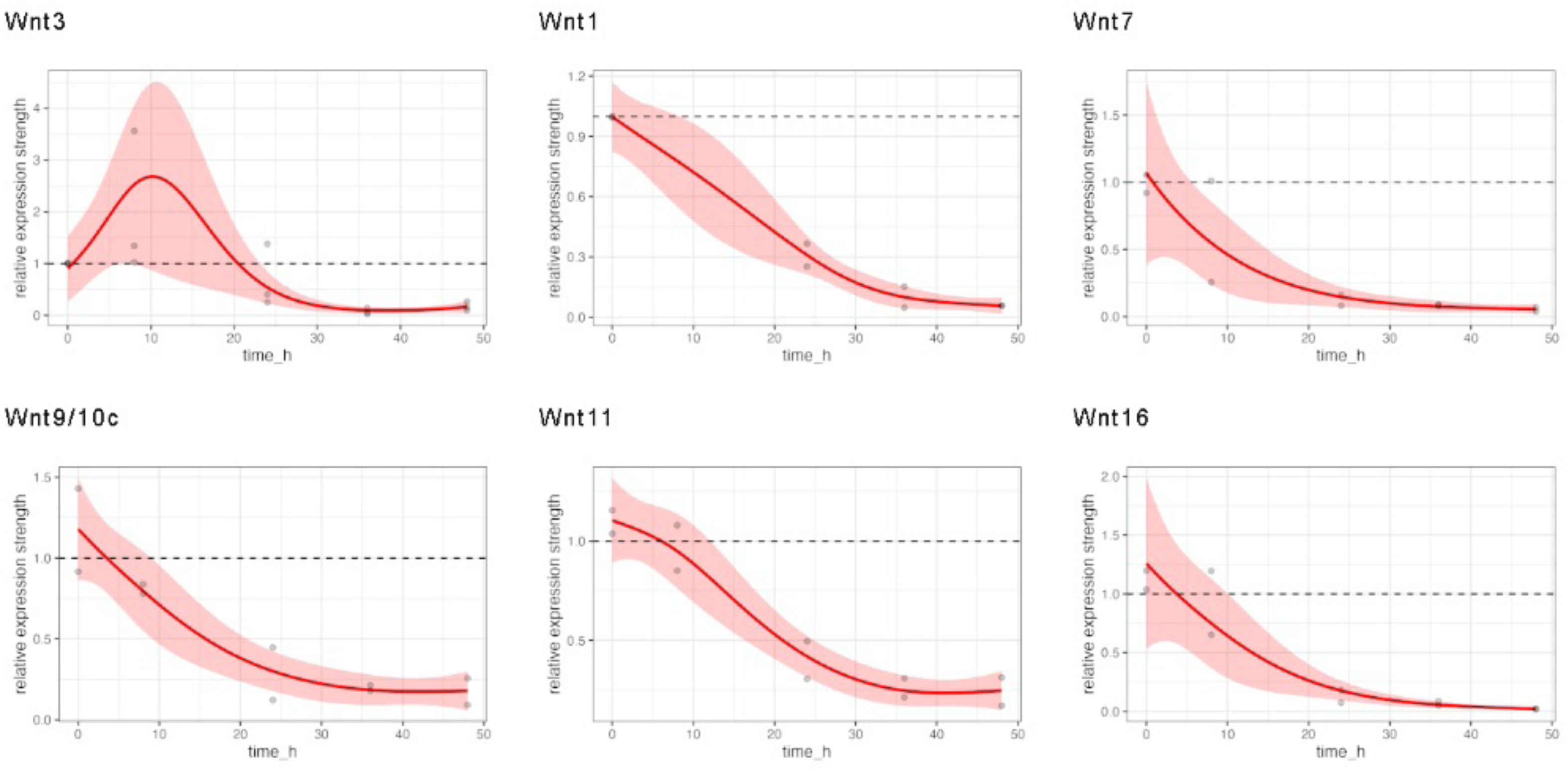
GAM-based visualization of relative gene expression dynamics of DAPT-treated regenerates. GAM-based visualisation of gene expression as measured by RT-qPCR in DAPT-treated animals relative to control animals (y-axis) depending on the time after head removal (x-axis). Grey points show raw data (quotients of mean values of DAPT-treated relative to DMSO-treated animals), the coloured lines show the smooth GAM-based estimates and colour-shaded areas are 95% confidence bands. Gene expression was followed for 48 hrs after head removal in DMSO control and DAPT. For time point 0 polyps were used immediately after head removal without any exposure to inhibitor or control medium. *HyHes, HyAlx, CnGsc, HySp5, HyKayak, HyBMP2/4* and *HyBMP5/8. HyWnt3* and *1, 7, 9/10c, 11, 16* during 48 hrs. Relative normalized expression was related to the housekeeping genes *GAPDH,* RPL13, EF1α and PPIB.

**Figure S4:**
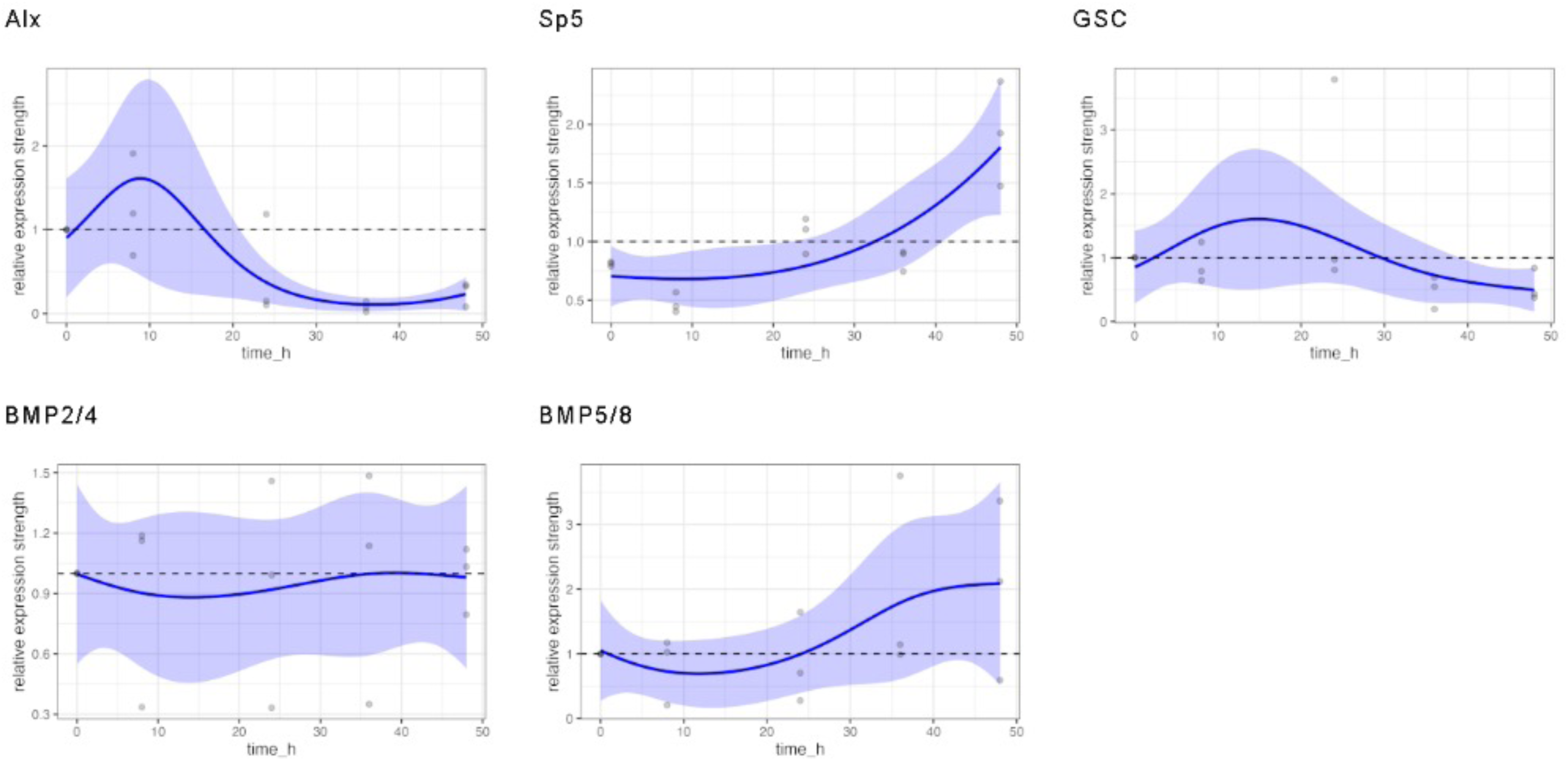
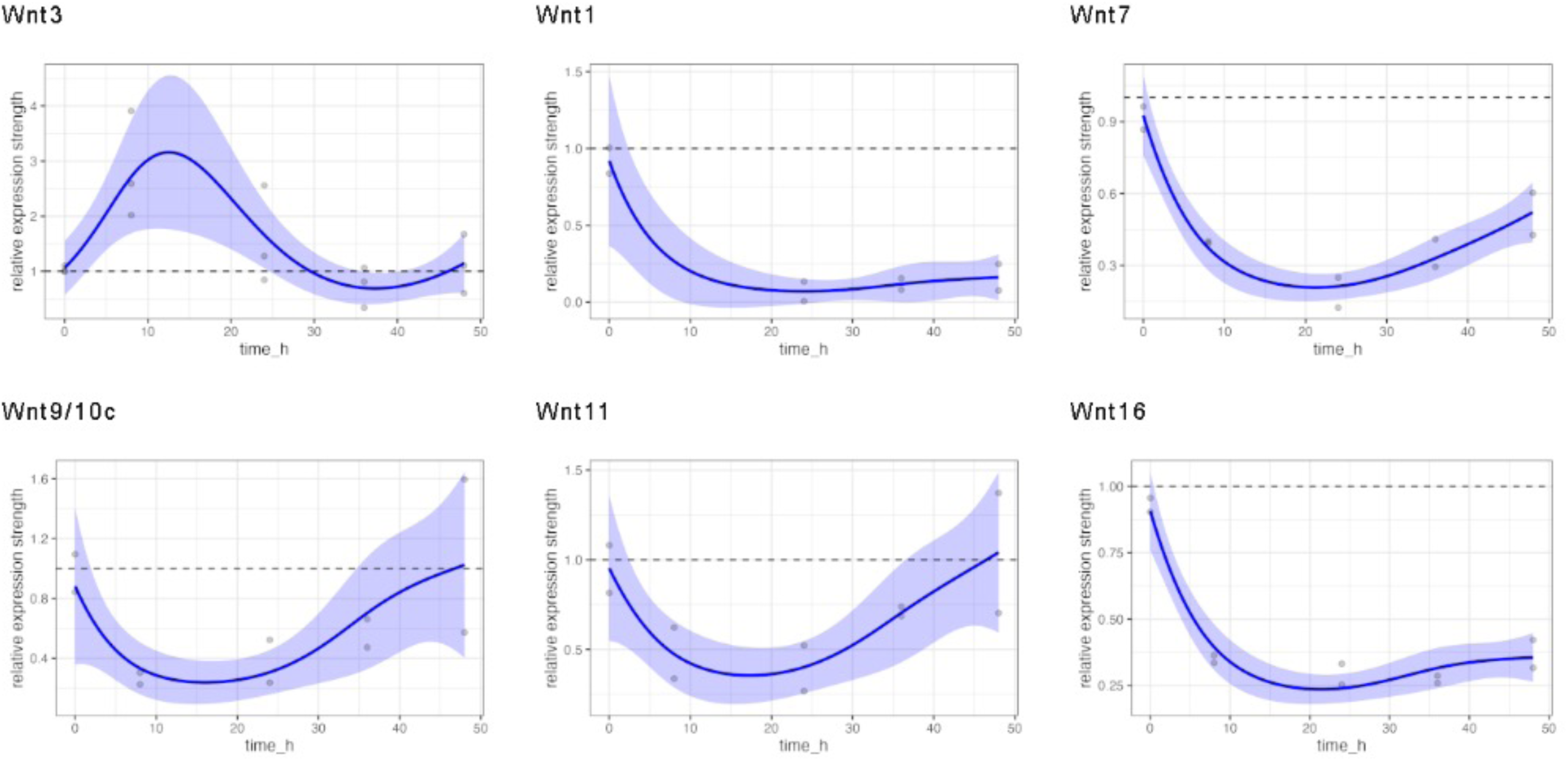
GAM-based visualization of relative gene expression dynamics of iCRT14-treated regenerates. GAM-based visualisation of gene expression as measured by RT-qPCR in iCRT14-treated animals relative to control animals (y-axis) depending on the time after head removal (x-axis). Grey points show raw data (quotients of mean values of iCRT14-treated relative to DMSO-treated animals), the coloured lines show the smooth GAM-based estimates and colour-shaded areas are 95% confidence bands. Gene expression was followed for 48 hrs after head removal in DMSO control and iCRT14. For time point 0 polyps were used immediately after head removal without any exposure to inhibitor or control medium. *HyHes, HyAlx, CnGsc, HySp5, HyKayak, HyBMP2/4* and *HyBMP5/8. HyWnt3* and *1, 7, 9/10c, 11, 16* during 48 hrs. Relative normalized expression was related to the housekeeping genes *GAPDH,* RPL13, EF1α and PPIB.

**Figure S5:**
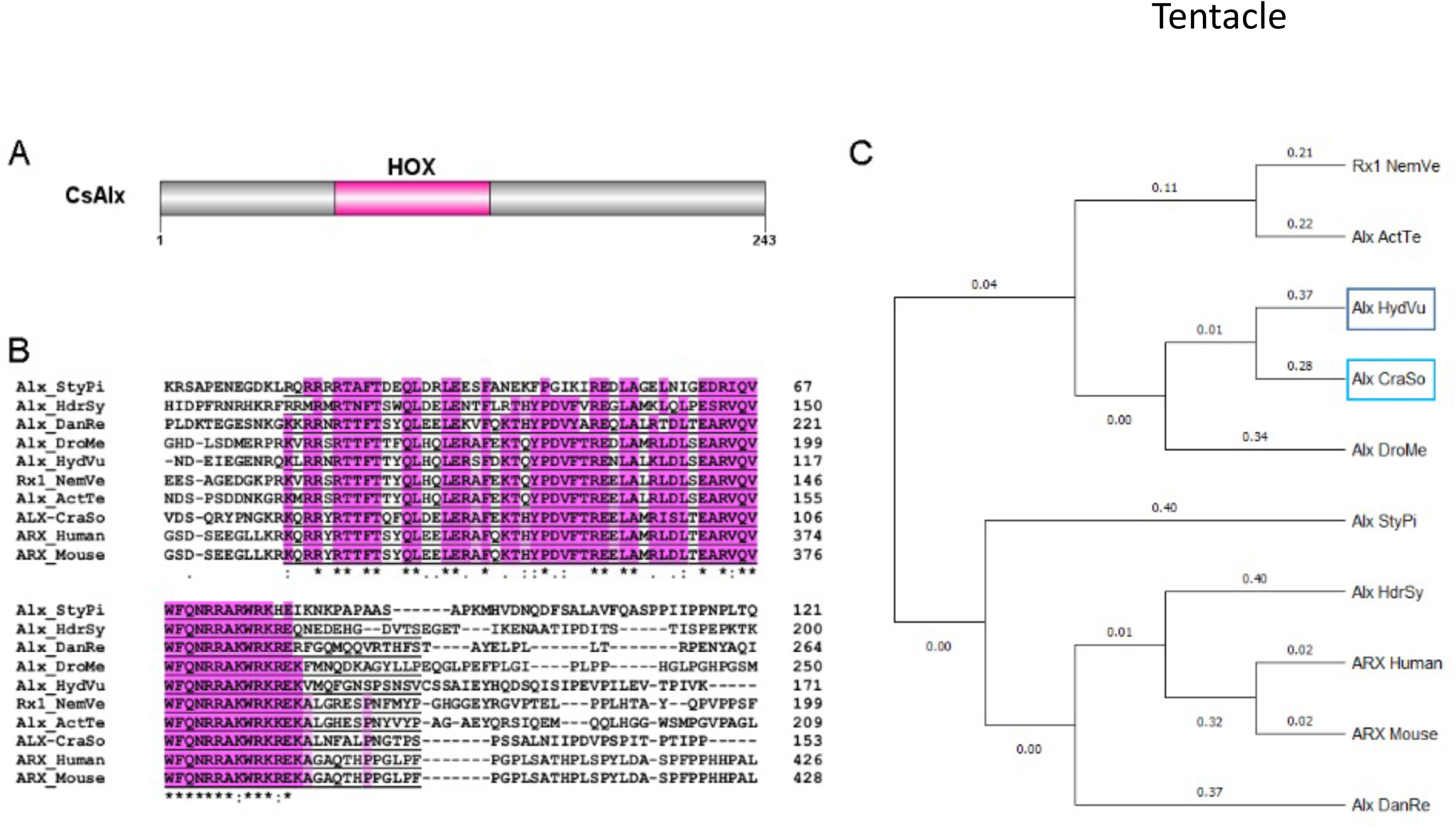
Craspedacusta CsAlx-gene identification and protein domain structure Sequence identification. **(A)** Schematic representation of the CsAlx protein structure (243 amino acids). The HOX domain with DNA binding function is located between amino acids 71 to 133. **(B)** Alignment of the protein sequences of Alx homologues from *Craspedacusta sowerbii* (CraSo), *Stylophora pistillata* (StyPi) – Acc#: PFX33415.1; *Hydractinia symbiolongicarpus* (HdrSy) – Acc#: XP_057291727.1; *Danio rerio* (DanRe) – Acc#: XP_001340966.1; *Drosophila melanogaster* (DroMe) – Acc#: NP_788420.1; *Hydra vulgaris* (HydVu) – Acc#: AAG03082.1; *Nematostella vectensis* (NemVe) – Acc#: XP_001634166.2*; Actnia tenebrosa* (ActTe) – Acc#: XP_031560466.1; Human - Acc#: NP_620689.1 and Mouse - Acc#: NP_001292869.1. The HOX domain is underlined, pink background indicates conserved amino acids involved in DNA-binding. **(C)** Phylogenetic tree based on the alignment of the 10 protein sequences affiliated to the aristaless family using MEGA software.

**Figure S6:**
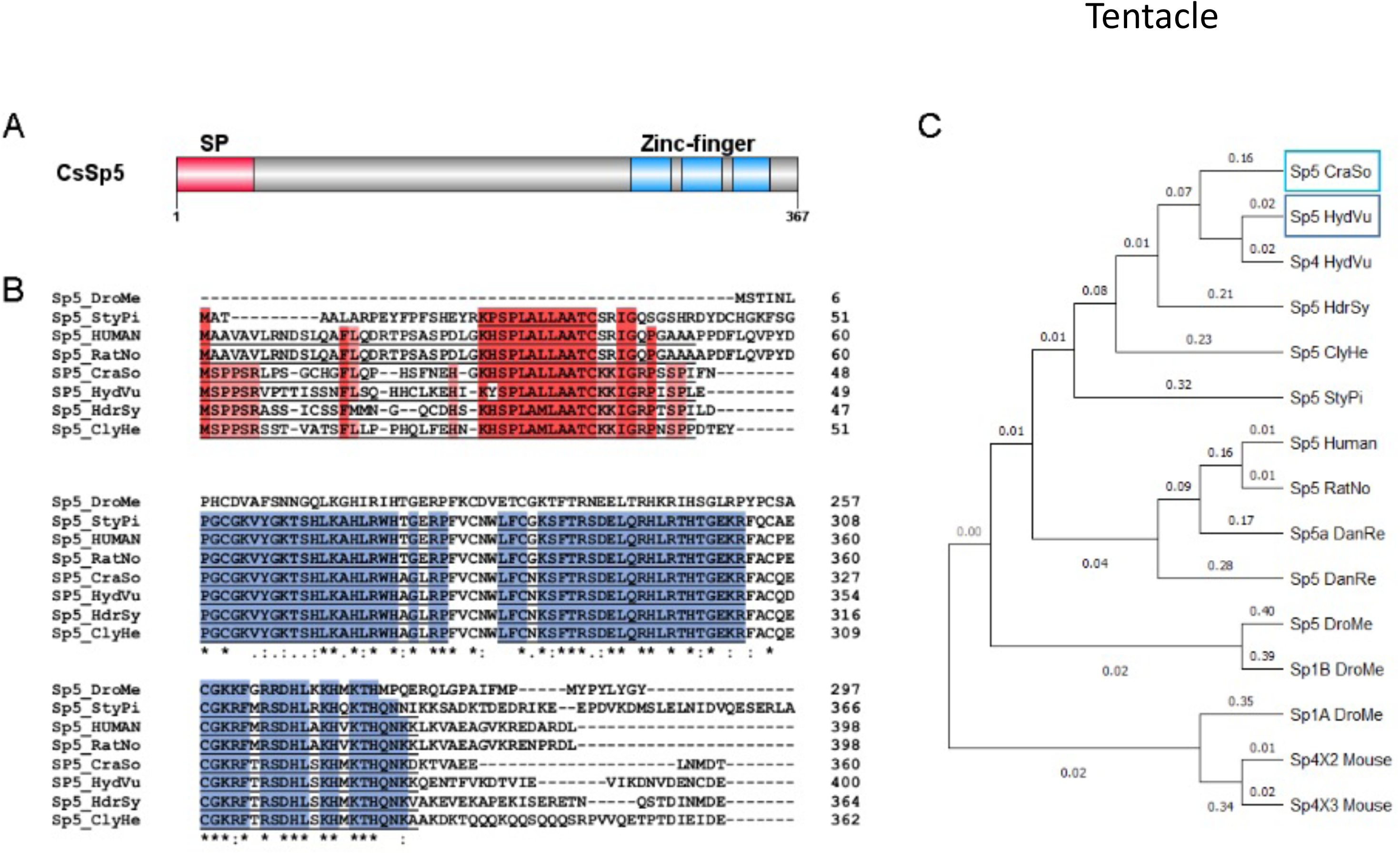
Craspedacusta CsSp5-gene identification and protein domain structure. **(A)** Schematic representation of the CsSp5 protein structure (367 amino acids). At the N-terminus the signal peptide (SP) for transclocating the protein is shown in red from amino acid 1 to 46. The three zinc-finger domains with DNA binding function are located at the C-terminus between amino acids 268-292, 298,322 and 328-350 shown in blue. **(B)** Alignment of the protein sequences of Sp5 homologues from *Craspedacusta sowerbii* (CraSo), *Drosophila melanogaster* (DroMe) – Acc#: NP_727360.1, NP_651232.1; *Stylophora pistillata* (StyPi) – Acc#: PFX28957.1; *Hydra vulgaris* (HydVu) – Acc#: AXP19710.1; *Hydractinia symbiolongicarpus* (HdrSy) – Acc#: XP_057304028.1; *Clytia hemisphaerica* (ClyHe) – Acc#: XP_057304028.1. The signal peptide is underlined, red background indicates conserved amino acids; Zinc-finger domains are underlined, blue background indicates conserved amino acids involved in DNA-binding. **(C)** Phylogenetic tree based on the alignment of the 15 protein sequences affiliated to the transcription factor Sp5 family using MEGA software. Human – Acc#: NP_001003845.1; *Rattus norvegicus* (RatNo) – Acc#: NP_001100022.1; *Danio rerio* (DanRe) – Acc#: NP_919352.1, NP_851304.2; Mouse – Acc#: XP_036013171.1, XP_036013172.1.

**Figure S7:**
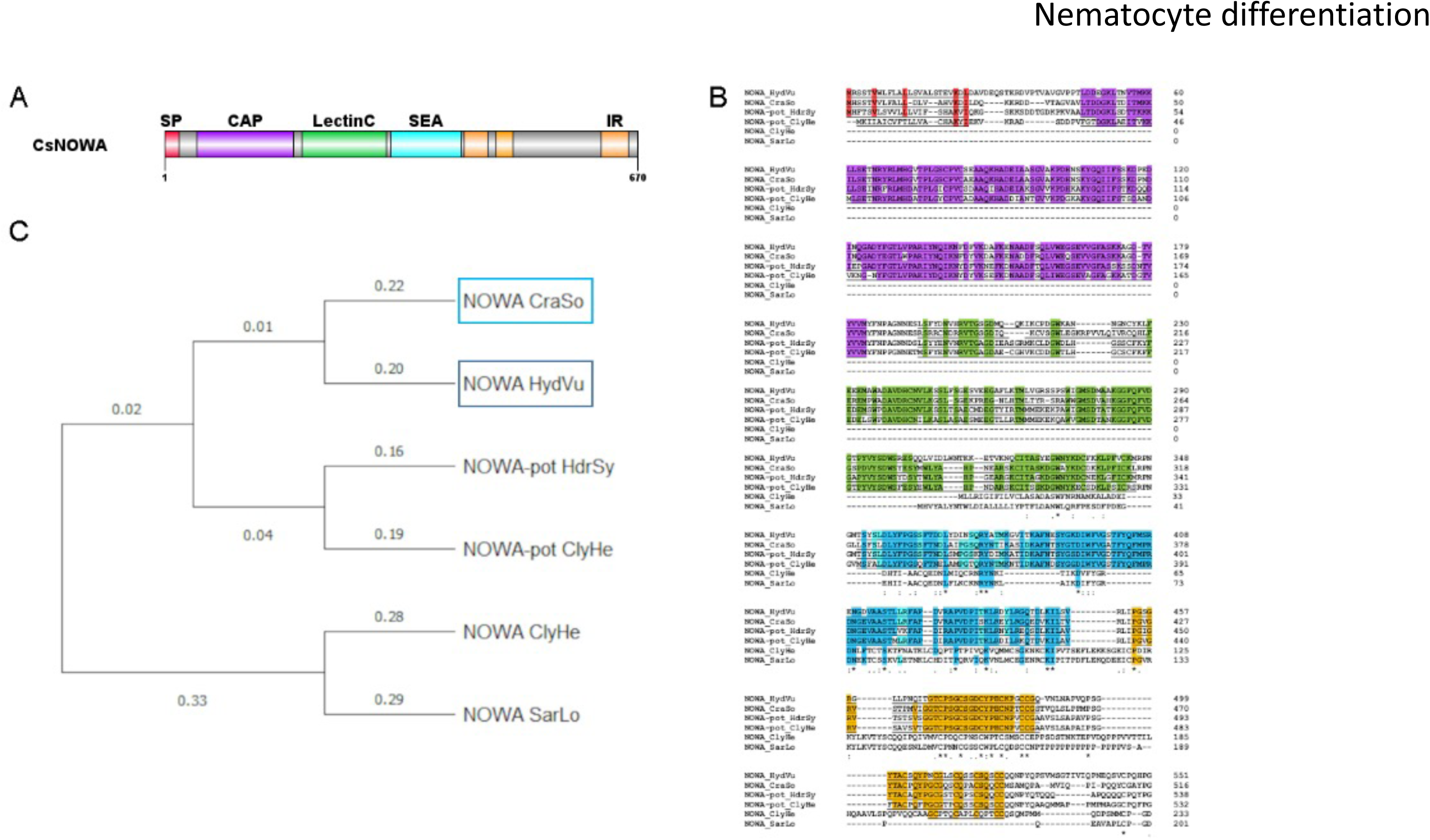
Craspedacusta CsNOWA-gene identification and protein domain structure. **(A)** Schematic representation of the CsNOWA protein structure (679 amino acids). At the N-terminus the signal peptide (SP) for transclocating the protein is shown in red from amino acid (aa) 1 to 20, followed by a CAP domain from 46-183 aa in violet, a carbohydrate recognition LectinC domain from 194-183aa shown in green and a SEA domain for membrane interaction shown in blue from 321-420 aa. The three internal repeats at the C-terminus are shown in orange between amino acids 424-457, 470-493 and 618-656. **(B)** Alignment of the protein sequences of NOWA homologues from *Craspedacusta sowerbii* (CraSo), *Hydra vulgaris* (HydVu) – Acc#: AAN52336.1; *Hydractinia symbiolongicarpus* (HdrSy) – Acc#: XP_057312482.1; *Clytia hemisphaerica* (ClyHe) – Acc#: XP_066935203.1 and precursor Acc#: ABY71251.1 and *Sarsia lovenii* (SarLo) – Acc#: WVX52206.1. The signal peptide is underlined, red background indicates conserved amino acids; CAP domains are underlined, violet background indicates conserved amino acids; LectinC domains are underlined, green background indicates conserved amino acids; SEA domains are underlined, blue background indicates conserved amino acids; Internal repeats are underlined, orange background indicates conserved amino acids. **(C)** Phylogenetic tree based on the alignment of the 6 protein sequences affiliated to the nematocyte producing antigen family using MEGA software.

**Figure S8:**
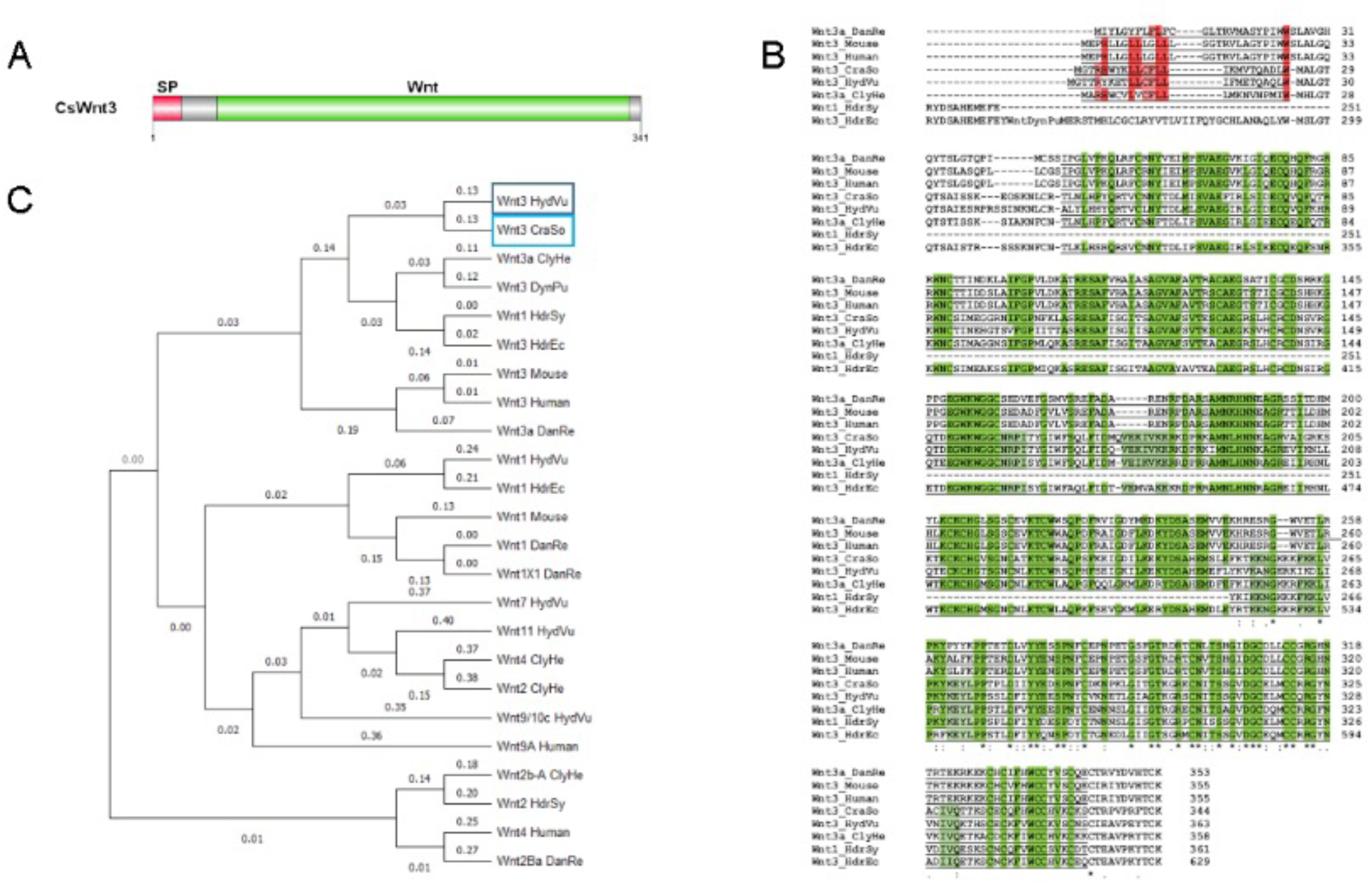
Craspedacusta CsWnt3-gene identification and protein domain structure. **(A)** Schematic representation of the CsWnt3 protein domain structure (341 amino acids). The N-terminal signal peptide (SP) is shown in red from amino acid (aa) 1 to 21, followed by the Wnt domain from 45-333 aa in green. **(B)** Alignment of the protein sequences of Wnt3 homologues from *Craspedacusta sowerbii* (CraSo), *Danio rerio* (DanRe) – Acc#: XP_005163717.1; Mouse – Acc#: NP_033547.1; Human – Acc#: NP_110380.1; *Hydra vulgaris* (HydVu) – Acc#: CDG70667.1; *Clytia hemisphaerica* (ClyHe) – Acc#: XP_066919214.1; *Hydractinia symbiolongicarpus* (HdrSy) – Acc#: XP_057304029.1 and *Hydractinia echinata* (HdrEc) – Acc#: CAK50826.1. The signal peptide is underlined, red background indicates conserved amino acids; Wnt domains are underlined, green background indicates conserved amino acids. **(C)** Phylogenetic tree based on the alignment of the 24 protein sequences affiliated to the Wnt3 family using MEGA software. *Hydra vulgaris:* Wnt1 - Acc#: BAH23782.1; Wnt7 – Acc#: BAH23781.1; Wnt11 – Acc#: BAH23776.1; *Clytia hemisphaerica* (ClyHe) – Acc#: XP_066919469.1, AFI99119.1, AFI99118.1; Dynamena pumila (DynPu) – Acc#: QBC65507.1; *Hydractinia symbiolongicarpus* (HdrSy) – Acc#: AIA10263.1; *Hydractinia echinata* (HdrEc) – Acc#: AIU99839.1; Mouse – Acc#: NP_067254.1; Human – Acc#: NP_110388.2, KAI4085194.1; *Danio rerio* (DanRe) – Acc#: NP_001188327.1, XP_005162280.1, NP_878296.1.

**Table S1:**
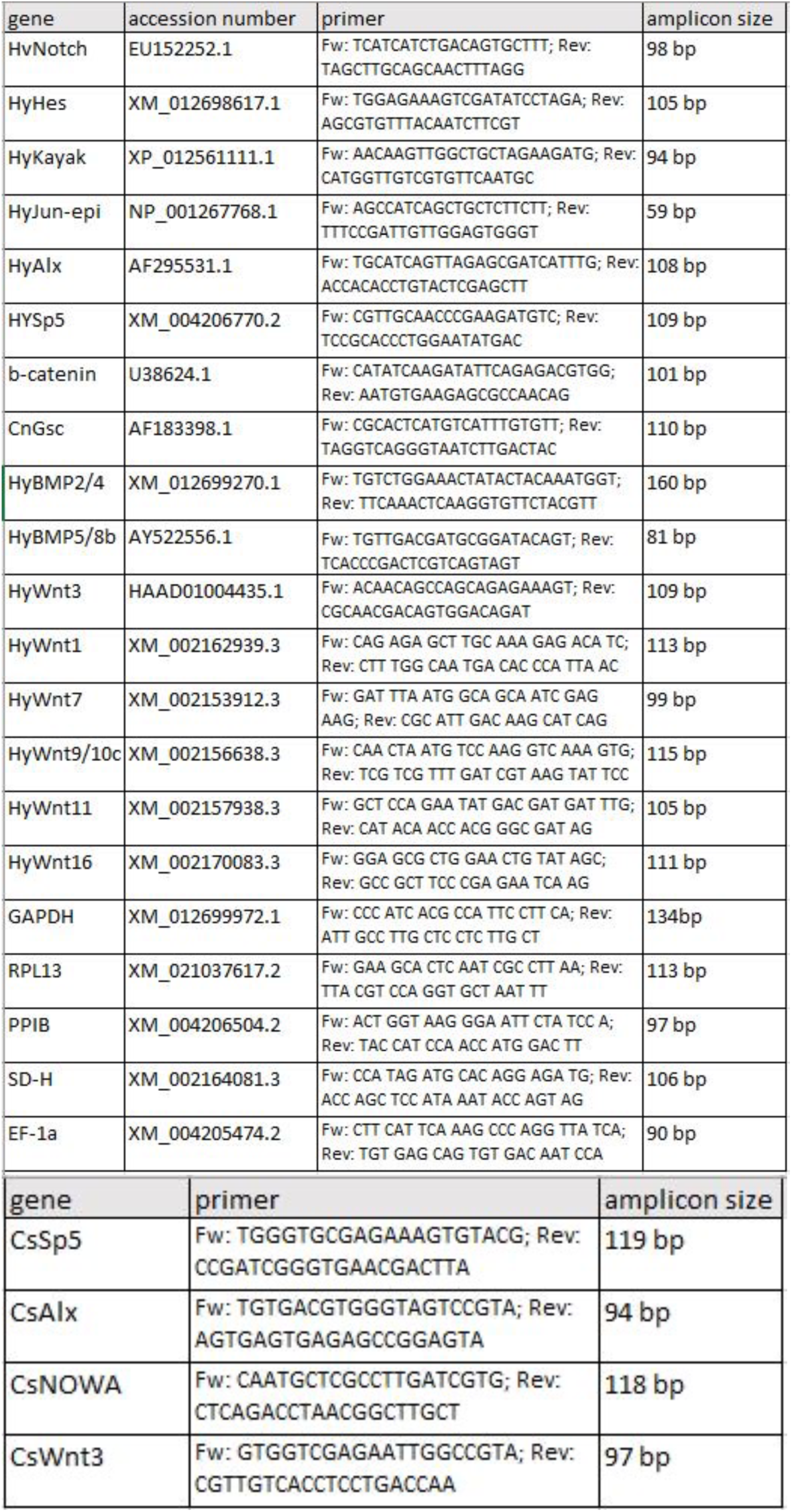
Primer list for qPCR.

